# FLT3-ITD signals for CEBPA and p53 proteolysis by the ubiquitin-proteosome pathway

**DOI:** 10.64898/2026.07.14.738455

**Authors:** Xiaorong Gu, Sudipta Biswas, Zeinab Albadry M. Zahran, Songa Bae, Ramesh Balusu, Jaroslaw P. Maciejewski, Babal K. Jha, Yogen Saunthararajah

## Abstract

Internal-tandem-duplication of the receptor tyrosine kinase *FLT3* (*FLT3-ITD*) generates ligand-independent signaling and is highly recurrent in acute myeloid leukemias (AMLs). One way signaling pathways can quickly influence cell fates is by phosphorylating key fate-determining proteins to trigger their proteolysis. We investigated the master transcription factor (MTF) driver of granulo-monocytic lineage-fates, CEBPA, for regulation by this mechanism because we found high CEBPA mRNA but little CEBPA protein in *FLT3-ITD* versus *FLT3*-wildtype AML cells, and inhibiting FLT3-ITD signaling with tyrosine kinase inhibitors (TKI) rapidly rescued CEBPA protein. Mass spectrometry analyses of CEBPA and its interactome demonstrated prominent interactions with major ubiquitin-proteosome pathway (UPP) components UHRF1 and USP7. TKI treatments decreased CEBPA and USP7 phosphorylations at serine 21 and serine 18 respectively alongside shifts in CEBPA interactions from degradative ubiquitin-ligase UHRF1 toward protective deubiquitinase USP7. The rescued CEBPA activated granulocytic-differentiation. Supporting that the serine-phosphorylations were ‘phospho-degrons’, UPP-inhibitors (bortezomib, MG132) increased phosphorylated and total CEBPA and USP7. The MTF regulator of apoptosis p53 is a known USP7 client, therefore, we also evaluated p53 status: TKIs and UPP-inhibitors stabilized USP7 and p53, triggering apoptosis in addition to granulocytic-differentiation specifically in *FLT3-ITD* but not *FLT3*-wildtype AML cells. UPP-inhibitors produced these consequences in TKI-resistant *FLT3-ITD* AML cells also. These data predicted genetic loss-of-function to *CEBPA* or *TP53* is redundant in the *FLT3-ITD* context, borne out by mutual exclusivity of the mutations in clinical series. In summary, FLT3-ITD signals for CEBPA and p53 proteolysis to block lineage-maturation and apoptosis, positioning UPP-inhibitors as therapeutic candidates acting downstream of TKIs.

**KEY POINTS:**

- The oncoprotein kinase FLT3-ITD signals for CEBPA and p53 proteolysis and hence suppresses lineage-differentiation and apoptosis
- Proteosome-inhibitors are candidate remedies to restore CEBPA and p53, acting downstream of presently used FLT3-ITD kinase inhibitors

**GRAPHICAL ABSTRACT:** 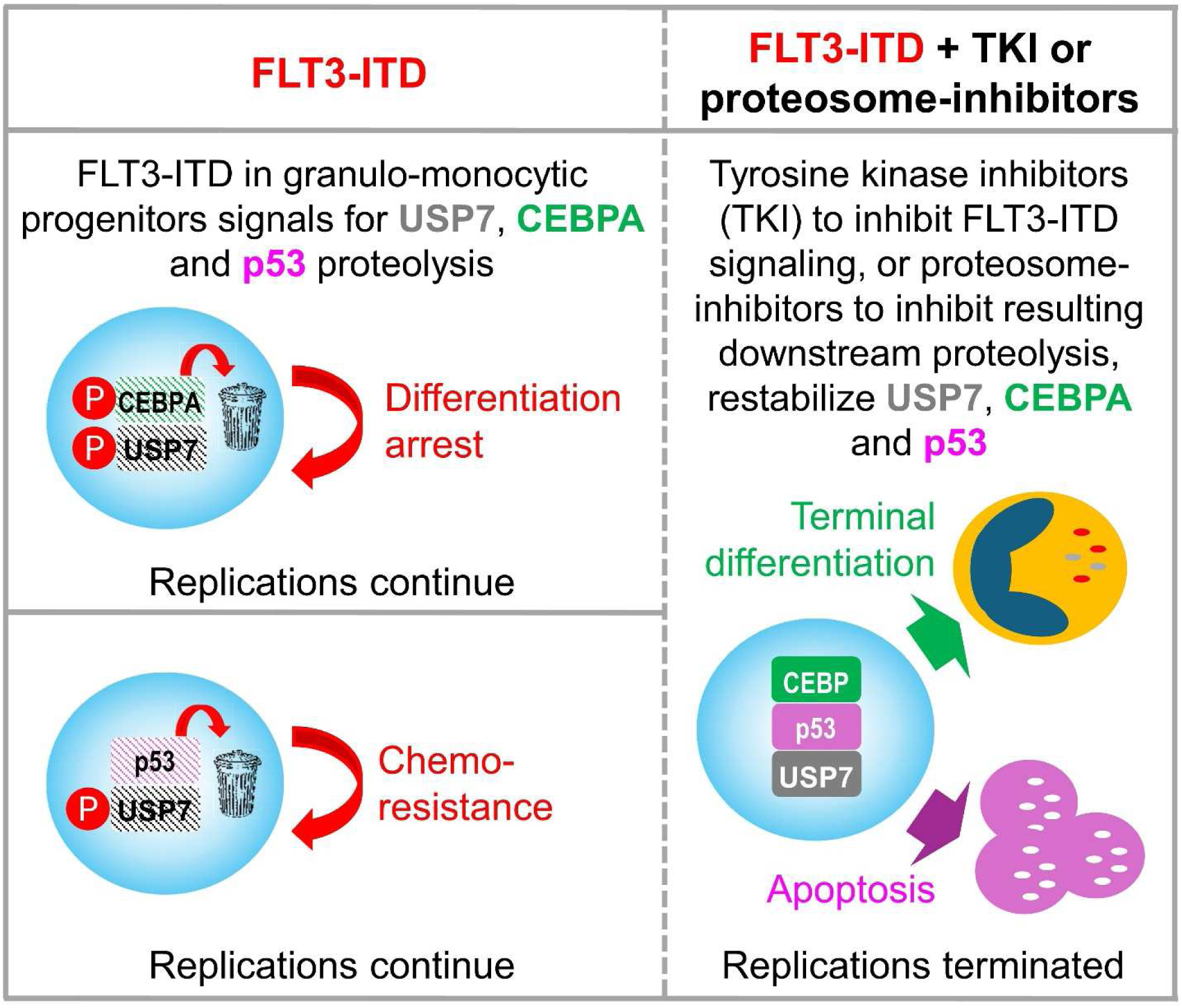

## INTRODUCTION

*Fms-related tyrosine kinase 3* (*FLT3*) is a class III receptor tyrosine kinase that is recurrently mutated in acute myeloid leukemias (AMLs) (∼30% of cases), most often as an internal-tandem-duplication (*FLT3-ITD*). FLT3-ITD is known to act differently from normal FLT3 in two key ways: first, it exercises constitutive, ligand-independent signaling activity(1, 2); second, FLT3-ITD is localized in an aberrant non-plasma membrane location and therefore signals from an origin and in a pattern also different from ligand-activated FLT3 (1–6). This aberrant signaling can be pharmacologically inhibited using several small molecule tyrosine kinase inhibitors (TKI) - midostaurin, gilteritinib, quizartinib, sunitinib, sorafenib – all of which have been used to treat *FLT3-ITD* AMLs, alone or combined with standard chemotherapy or hypomethylating agents (HMAs) (reviewed in (7)). Nevertheless, five-year survivals for patients with *FLT3-ITD* AMLs are <30% (8). A better understanding of the downstream molecular consequences of FLT3-ITD kinase signaling, that transform normal into malignant myelopoiesis, can hopefully point to new treatment modalities.

In normal myelopoiesis, granulocyte and monocyte precursor cells (granulo-monocyte progenitors, GMPs) divide frequently, e.g., every 1-2 days, to propel exponential growth kinetics as is needed to replace tens of billions of granulocytes and monocytes lost to wear and tear each day. This exponential growth is self-limiting because with each division, daughter cells activate lineage-specific genes, arriving after several rounds of division at final cell identities wherein replications are halted to focus on specialist granulocyte or monocyte roles. FLT3-ITD is known to disrupt this granulo-monocytic lineage-differentiation process specifically: (i) *FLT3-ITD* originates in GMPs and is not found in hematopoietic stem cells (HSC) that sit upstream in the hematopoietic hierarchy (9–12); (ii) the precursor cells stalled in lineage-differentiation and that continue to replicate and accumulate in *FLT3-ITD* AMLs – myeloblasts - phenocopy GMPs (12–14); (iii) increasing the dose of *FLT3-ITD* via copy neutral loss-of-heterozygosity decreased granulo-monocytic lineage-differentiation and increased clonal expansion in parallel (15); (iv) *FLT3-ITD* is highly recurrent in AMLs with granulo-monocytic lineage-features including acute promyelocytic leukemias (∼40% of cases) but is absent from AMLs with erythroid or megakaryocyte lineage-features; and (v) *FLT3-ITD* is very rare in the myeloid neoplasm myelodysplastic syndromes (MDS) without myeloblasts, but is found in MDS with myeloblasts (16, 17), and is over-represented in AMLs with the highest myeloblast counts (hyperleukocytic AMLs) (18).

Altogether, therefore, FLT3-ITD has been linked to decoupling onward lineage-differentiation from replications in GMPs to generate myeloblasts. Driving this lineage-differentiation normally is a combinatorial code of master transcription factors (MTFs)(19) which includes CCAAT enhancer binding protein alpha (CEBPA) - CEBPA is indispensable to granulopoiesis and contributes to monopoiesis (reviewed in(20)). CEBPA has been previously connected to FLT3-ITD signaling: (i) Inhibiting FLT3-ITD signaling in an AML cell line (MV4-11) using a TKI decreased a post-translational modification of CEBPA (serine 21 phosphorylation, S21-P) and released terminal granulocytic lineage-differentiation (21); (ii) knocking-down CEBPA in another *FLT3-ITD* AML cell line (MOLM14) prevented the terminal granulocytic-differentiation response to a TKI (22); and (iii) *FLT3-ITD* introduction into murine granulo-monocytic precursor cells (32D) prevented the granulocytic differentiation response to granulocyte-colony stimulating factor (G-CSF) and the investigators proposed the mechanism for this was less transcription of *Cebpa* (23, 24).

Here, in examining connections between FLT3-ITD and CEBPA further, we observed high CEBPA mRNA but little CEBPA protein in *FLT3-ITD* AML cells. Regulation of MTFs via their continuous proteolysis, that can be modulated by signaling inputs, is a known motif in biology (25). For example, in normal proliferating myeloid progenitors, *TP53* that encodes for p53, the MTF regulator of apoptosis, is continuously transcribed and p53 is instead regulated at the protein level (25–28), including by USP7, a key deubiquitinase in the ubiquitin-proteosome pathway (UPP) (29–32). Regulation of MTFs by the proteosome in this way circumvents delays inherent to transcription-translation, enabling rapidly dividing cells to mount immediate MTF-driven fate and function responses to appropriate stimuli (25–28, 33). We found that FLT3-ITD signaling hijacks this motif to continuously degrade the MTFs CEBPA and p53 that normally regulate myeloid progenitor growth. A translational implication is that UPP-inhibitors, presently used to treat other cancers (34, 35), could be candidate rational therapeutics to remedy the FLT3-ITD transforming pathway at points further downstream than engaged by TKI.

## METHODS

### Sources of cell lines and animals

MV4-11 and THP1 AML cell lines were purchased from ATCC (Manassas, Virginia). MOLM13 and OCI-AML3 AML cell lines were purchased from DSMZ (Braunschweig, Germany). The cell lines were additionally authenticated (Labcorp Cell Line Authentication, Burlington, NC).

### Key chemicals, reagents and antibodies used

*MedChem Express*

Sunitinib, HY-10255A; Gilteritinib, HY-12432; Bortezomib, HY-10227; MG-132, HY-13259. *Sigma Aldrich*: Benzonase Nuclease, E1014; Protease inhibitor cocktail, P8340; Phosphatase inhibitor cocktail 3, P0044; Phosphatase inhibitor cocktail 2, P5726. BioRad: PVDF membranes, 1620177; Precision Plus Protein Dual Color Standards, 1610374; Anti-phospho USP7 (ser18), ABC225. *Invitrogen*: NuPAGE MOPS SDS Running Buffer, NP0001; NuPAGE Transfer Buffer, NP0006; Goat anti-Rabbit IgG, A32735; Goat anti-Mouse IgG, A32730; Goat anti-Rabbit IgG, A21038; Goat anti-Mouse IgG, A21036. *Santa Cruz Biotech*: Protein A/G PLUS-Agarose, SC-2003; Normal mouse IgG, SC-2025; Anti-CEBPA (G-10), SC-166258. *Cell Signaling*: Anti-Phospho-CEBPA (ser21), 2841S; Anti-HAUSP (USP7) (D17C6), 4833S; Anti-p53, 9282S.

### Immunofluorescence microscopy

Cells were cyto-spinned on to glass slides and fixed in cold methanol for 30min at -20°C. The fixed cells were blocked in 5% bovine serum albumin for 1 hour at room temperature, then incubated overnight at 4°C with the primary antibodies CEBPA (1:200, Cat# sc-166258, Santa Cruz Biotechnology) and CEBPA S21-P (1:200, Cat# 2841, Cell Signaling). Cells were washed in 1xPBS with 0.1% Tween 20 followed by incubation with secondary antibodies, Alexa Fluor 488 Goat Anti-Mouse (1:250, A1101, Invitrogen) or Alexa Fluor 568 Goat Anti-Rabbit IgG (1:250, A11036, Invitrogen), for one hour at room temperature in the dark. Cells were washed in 0.1% Tween 20 and 1X PBS. Nuclei of cells were stained with DAPI (4′,6-diamidino-2-phenylindole) mounting medium (Vector, H-1200). Images were taken with Leica DM RBE microscope, connected to CRI Nuance multispectral imaging camera, running Nuance version 3.0.2 software (PerkinElmer). Confirmatory images were acquired using Leica SP8 inverted confocal microscope (Leica Microsystems, GmbH, Wetzlar, Germany) running Leica Application Suite X software.

### Cell fractionation, nuclear protein extraction, covalent binding of antibodies to protein A/G beads, immunoprecipitation, 1D SDS-polyacrylamide gel electrophoresis and Western blot analysis

As we have previously described (36, 37).

### NanoLC-ESI-LTQ-Orbitrap MS/MS

Immunoprecipitation products were subjected to SDS-polyacrylamide gel electrophoresis and stained with colloidal Coomassie Blue (Gel Code Blue, Pierce Chemical). Gel slices were excised from the top to the bottom of the lane; proteins were reduced with dithiothreitol (Sigma-Aldrich, D0632, 10mM), alkylated with iodoacetamide (Sigma-Aldrich, I1149, 55mM), and digested in situ with trypsin. Peptides were extracted from gel pieces 3 times using 60% acetonitrile and 0.1% formic acid/water. The dried tryptic peptide mixture was redissolved in 20 μL of 1% formic acid for mass spectrometric analysis. Tryptic peptide mixtures were analyzed by on-line LC-coupled tandem mass spectrometry (LC-MS/MS) on an Orbitrap mass spectrometer (Theomo Fisher Scientific).

### Database Search and Data Validation

Mascot Daemon software (version 2.3.2; Matrix Science, London, UK) was used to perform database searches, using the Extract_msn.exe macro provided with Xcalibur (version 2.0 SR2; Thermo Fisher Scientific) to generate peaklists. The following parameters were set for creation of the peaklists: parent ions in the mass range 400–4500, no grouping of MS/MS scans, and threshold at 1000. A peaklist was created for each analyzed fraction (i.e., gel slice), and individual Mascot (version 2.3.01) searches were performed for each fraction. The data were searched against Homo sapiens entries in Uniprot protein database (Feb 2018 release; 20,316 total sequences). Carbamidomethylation of cysteines was set as a fixed modification, and oxidation of methionine was set as a variable modification. Specificity of trypsin digestion was set for cleavage after Lys or Arg, and two missed trypsin cleavage sites were allowed. The mass tolerances in MS and MS/MS were set to 10 ppm and 0.6 Da, respectively, and the instrument setting was specified as “ESI-Trap.” To calculate the false discovery rate (FDR), the search was performed using the “decoy” option in Mascot. The spectral FDR and protein FDR are 0.35±0.17 % and 4.28±1.99 % respectively. A minimum Mascot ion score of 25 and peptide rank 1 was used for automatically accepting all peptide MS/MS spectra.

### Label free relative protein quantitation (LFQ)

Relative protein quantification was performed using spectral count-based LFQ. For each biological sample, data from the individual gel slices were combined. LC-MS/MS data files were imported into Agilent MassHunter Mass Profiler Professional (MPP) software. Identified proteins, along with their label-free quantification (LFQ) values, were filtered based on their presence in at least 75% of replicates within each sample group and a minimum abundance threshold to reduce technical noise. The proteomic data has been uploaded into ProteomeXchange (accession number pending).

### Giemsa staining of cells

Cytospins of cells from bone marrow or peripheral blood were fixed for 2 minutes in methanol, air-dried, and stained for 20 minutes with filtered modified solution of Giemsa stain (Sigma Aldrich, Cat # 48900, St Louis, MO), diluted (1:5) with buffer solution pH6.5, rinsed with distilled water, air-dried and examined using low and high magnifications with a Leica DMR microscope (Leica Microsystems, Wetzlar GmbH, Germany) connected to Nuance multispectral imaging system FX using Nuance version 3.0.2 software (PerknElmer, Inc., Hopkinton, MA).

### Flow Cytometry Analyses for CD11b

As we have previously described (36, 38). Antibodies used were anti-human CD11b (clone: M1/70, cat. no. 101206, Biolegend, 1:100).

### Apoptosis detection

Apoptosis was detected by Annexin-V and propidium iodide (PI) co-staining using the APOAF commercial kit (Sigma). Cells (5×10^5^) were washed and incubated for 30 minutes with FITC-conjugated Annexin-V at room temperature. Cells were then resuspended in 400 mL of binding buffer containing PI and immediately analyzed by flow cytometry.

### RNA isolation, reverse transcription (RT) and real-time PCR

Total RNA from cultured cells was isolated using RNeasy Mini Kit (Qiagen, Cat# 74104) according to the manufacturer’s instruction. The cDNA was then synthesized from total RNA using the iScript cDNA synthesis Kit (BioRad, Cat# 1708891). Quantitative gene expression levels were detected using real-time PCR with the ABI PRISM 7500 Fast Sequence Detection System and SYBR Advantage qPCR Premix (Clontech, 639676) according to the manufacturer’s instructions. Primers for all genes analyzed were purchased from Integrated DNA Technologies (primer sequences are provided in supplementary material). Relative expression values (RQ values) were calculated by the delta-delta C_T_ method with beta-actin or GAPDH as the internal control.

### Bioinformatic and statistical analysis

#### Identification of CEBPA target-genes that are normally upregulated with granulo-monocytic differentiation

FastQ files from SRA for CEBPA and IgG control chromatin-immunoprecipitation sequencing (ChIP-seq) in GMPs (SRR935459, SRR636721 respectively) were downloaded and processed using the UseGalaxy suite of tools (39). Aligned ChIP-Seq reads were imported, analyzed, and visualized using Easeq(40), including the EASEQ peak-finding tool(40). Values were normalized to reads per million per 1kbp, and genes with Cebpa binding peaks within 500 base pairs of the gene transcription start sites that were >3-fold higher than observed with the IP IgG control were identified. The subset of these genes which were upregulated during normal myeloid ontogeny into granulocytes and monocytes (BloodSpot (41)), and which had a statistically significant correlation in their expression with CEBPA expression in the same cells (Pearson correlation coefficient, p<0.005, 2-tailed) were identified as CEBPA target genes activated normally during lineage maturation of GMPs. The Comparative Marker Selection tool in Morpheus (https://software.broadinstitute.org/morpheus/, Broad Institute, MIT, USA (42), an algorithm for identifying genes that discriminate between classes of samples, 1000 permutations test, p-value ≤0.02, was used to identify the genes differentially expressed during normal myeloid ontogeny.

#### Identifying CEBPA interacting proteins

IPs of endogenous CEBPA were performed with CEBPA antibody or control IgG isotype antibody. Total spectra counts (x + 1) for individual proteins were compared between the IP-CEBPA and IP-IgG control using an unpaired 2-sided t-test. A Benjamini-Hochberg adjusted p-value <0.05 was considered significant. For comparisons between *FLT3-ITD* vs *FLT3*-wildtype AML cells, spectral count data was normalized by total spectral counts of the bait protein (CEBPA or USP7) in each sample.

Wilcoxon rank sum, Mann Whitney, and t tests were 2-sided and performed at the 0.05 significance level or lower (Bonferroni or Benjamini-Hochberg corrections were applied for instances of multiple parallel testing). Standard deviations (SD) and inter-quartile ranges (IQR) for each set of measurements were calculated and represented as y-axis error bars on each graph. Graph Prism (GraphPad, San Diego, CA) or SAS statistical software (SAS Institute Inc., Cary, NC) was used to perform statistical analysis including correlation analyses.

## RESULTS

### CEBPA mRNA and protein amounts in *FLT3-ITD* versus other AML cells

We had previously observed that primary AML cells activate *CEBPA* to the extent that CEBPA mRNA levels are multi-log-fold higher than seen in normal HSCs, and frequently exceed levels in normal GMP or terminally-differentiated granulocytes and monocytes (37). We parsed this data to better understand connections between FLT3-ITD and CEBPA, and observed that CEBPA mRNA was similarly elevated in *FLT3*-wildtype (WT) (n=412) and *FLT3-ITD* (n=137) primary AML cells (gene expression by RNA-sequencing, BEAT AML public data (43)) (**Figure 1A**). Then, to investigate downstream actions of the activated *CEBPA*, we examined expression of CEBPA target-genes in the cells, that is, genes bound by Cebpa in their proximal promoters (44) and activated during normal granulo-monocytic lineage-differentiation (n=347, **Table S1**). For these analyses we included as an additional control primary AML cells containing bi-allelic *CEBPA* mutations (*CEBPA*^dm^ - double-mutant) – by truncating CEBPA and/or preventing its DNA-binding, these mutations are expected to attenuate activation of CEBPA target-genes (45). CEBPA target-genes were suppressed in *CEBPA*^dm^ vs *FLT3*-wildtype primary AML cells as expected (**Figure 1B**). Unexpectedly, however, CEBPA target-genes were also suppressed in *FLT3-ITD* primary AML cells to the low levels seen in the *CEBPA*^dm^ cells (**Figure 1B**). These results were seen also in a separate clinical cohort of primary AML samples (gene expression by RNA-sequencing, TCGA AML public data)(**Figure 1C**, **D**). These data implied that *CEBPA*^dm^ is redundant in the *FLT3-ITD* context: accordingly, *CEBPA*^dm^ was mutually exclusive with *FLT3-ITD* in primary AML cohorts, even though both sets of mutations are highly recurrent (**Figure S1**).

**Figure 1.**
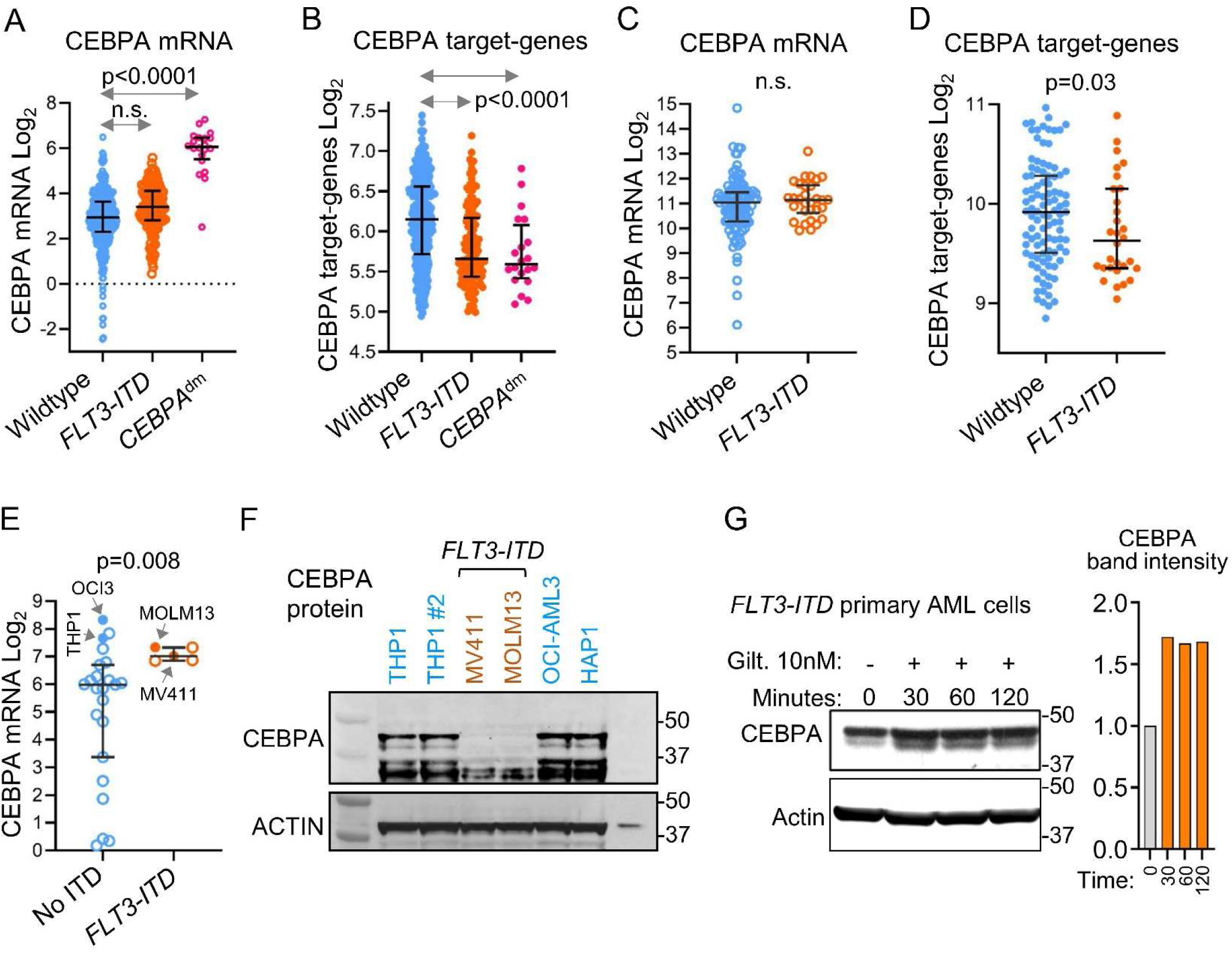
Attenuated CEBPA target-gene activation in *FLT3-ITD* AML cells is explained by less CEBPA protein. **A) CEBPA mRNA levels in *FLT3*-wildtype (WT) (n=412), *FLT3-ITD* (n=137), and *CEBPA*^dm^ (n=20) primary AML cells.** AMLs containing CEBPA mutations that truncate and/or prevent DNA binding by CEBPA (*CEBPA*^dm^) were included as additional controls expected to attenuate CEBPA target-gene activation. Gene expression by RNA-sequencing, conditional quantile normalization of gene level counts was described previously(43) (BEAT AML public data; acute promyelocytic leukemia samples excluded). Median ± IQR. P-value Mann-Whitney test, 2-sided. **B) Activation of CEBPA target-genes in *FLT3*-WT (n=412), *FLT3-ITD* (n=137), and *CEBPA*^dm^ (n=20) primary AML cells.** AMLs containing CEBPA mutations that truncate and/or prevent DNA binding by CEBPA (*CEBPA*^dm^) were included as additional controls expected to attenuate CEBPA target-gene activation. Average expression (by RNA-sequencing as per panel A) of 347 Cebpa target-genes is plotted. Cebpa target-genes contained Cebpa peaks in their proximal promoters(44) and were activated during differentiation of GMPs into granulocytes and monocytes (**Table S1**). Median ± IQR. P-value Mann-Whitney test, 2-sided. **C) CEBPA mRNA levels in a separate clinical cohort of *FLT3*-wildtype (WT) (n=101) and *FLT3-ITD* (n=30) primary AML cells.** mRNA levels measured by RNA-sequencing, log_2_ transformed RSEM normalized count (TCGA public data. Samples containing *CEBPA* mutations, and acute promyelocytic leukemia samples were excluded from the analyses). Median ± IQR. P-value Mann-Whitney test, 2-sided. **D) Activation of CEBPA target-genes in this separate clinical cohort.** Average expression (by RNA-sequencing as per panel C) of 347 Cebpa target-genes (Table S1) is plotted. Median ± IQR. P-value Mann-Whitney test, 2-sided. **E) We identified *FLT3*-wildtype and *FLT3-ITD* AML cell lines expressing similar levels of CEBPA mRNA.** mRNA levels in granulo-monocytic AML cell lines characterized for *FLT3-ITD* presence or absence, Table S2(69, 70) (RNA-sequencing, RPKM, Cancer Cell Line Encylopedia). Median ± IQR. P-values Mann-Whitney test, 2- sided. **F) The *FLT3-ITD* AML cells (MV4-11, MOLM13) contained little CEBPA protein compared to *FLT3*-WT AML cells (THP1, OCI-AML3, HAP1).** Western blots. **G) Inhibiting FLT3-ITD signaling with gilteritinib rapidly upregulated CEBPA protein in *FLT3-ITD* primary AML cells.** Primary AML cells from a patient with *FLT3-ITD* AML treated with DMSO vs Gilteritinib 10nM. Graphs = fold-change relative to DMSO treatment of CEBPA band intensity normalized to Actin. Western blots.

Since diminished activation of CEBPA target-genes in *FLT3-ITD* AML cells was not explained by diminished CEBPA mRNA (23, 24), we examined if FLT3-ITD signaling diminished CEBPA protein. We identified *FLT3*-wildtype (THP1, OCI-AML3, HAP1) or *FLT3-ITD* (MOLM13, MV4-11) AML cell lines that contained similar amounts of CEBPA mRNA (**Figure 1E**, **Table S2**) and examined their CEBPA protein content: CEBPA protein was abundant in the *FLT3*-wildtype- but markedly diminished in the *FLT3-ITD*-AML cells despite their similar amounts of CEBPA mRNA (**Figure 1F**). To extend this data to primary *FLT3-ITD* AML cells, we inhibited FLT3-ITD signaling with the TKI gilteritinib 10nM: CEBPA protein was upregulated within minutes (**Figure 1G**).

### CEBPA interacts with major UPP components, under control by FLT3-ITD signaling

The speed at which CEBPA protein was rescued by the TKI treatment of the primary *FLT3-ITD* AML cells suggested regulated engagement of CEBPA protein by proteosome pathways. We therefore studied endogenous steady-state CEBPA protein and its interactions by immunoprecipitation-liquid chromatography tandem mass spectrometry (IP-LCMS/MS) as we have described previously for other MTFs (36, 37, 46, 47). Since *FLT3-ITD* AML cells contain less CEBPA protein (**Figure 1G**), we used 3-fold more of these cells per extraction (MOLM13 ∼150 million cells each) than *FLT3*-wildtype cells (OCI3, THP1, ∼50 million each) - spectral count data confirmed enrichment for this bait protein (**Table S3**). Using IP with IgG-isotype as a control, we identified 127 proteins interacting with CEBPA (**Table S3**). Of these 127 proteins, 15 pulled-down with abundancies paralleling the CEBPA bait protein, suggesting the highest probability interactions (**Table S3**). These included major UPP components ubiquitin-ligase UHRF1 and deubiquitinase USP7 (**Figure 2A, Table S3**), that were moreover ∼4-fold more represented in the CEBPA interactome in the *FLT3-ITD* than *FLT3*-wildtype AML cells (Benjamini-Hochberg adjusted p-value <0.05) (**Figure 2A, Table S4**).

**Figure 2.**
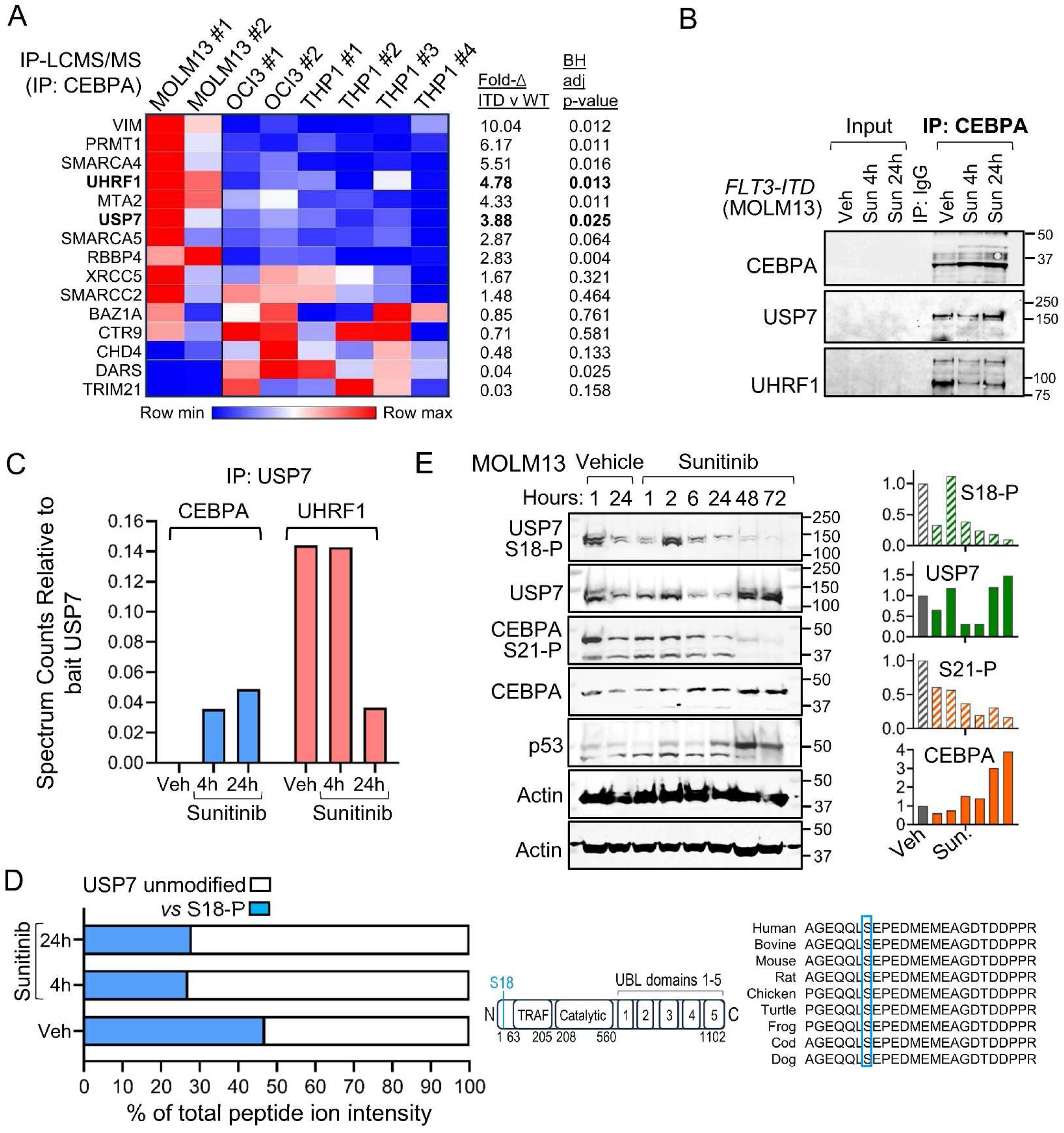
CEBPA interacts with key ubiquitin-proteosome pathway (UPP) components, influenced by FLT3-ITD signaling. **A) Protein-protein interactions of CEBPA in *FLT3-ITD* (MOLM13)** *vs FLT3*-wildtype **(WT) AML cells (THP1, OCI-AML3).** Endogenous CEBPA was immunoprecipitated (IP) and interacting proteins identified by LC-MS/MS and Proteome Discovery and Mass Profiler Professional (MPP) software. Interacting proteins (127 proteins, Table S3) had total spectra counts significantly increased (Benjamini-Hochberg adjusted p-value <0.05) *vs* IP with IgG isotype control antibody. Fifteen of the 127 proteins had abundancies similar to CEBPA bait-protein, including UHRF1 and USP7 which were significantly enriched in the CEBPA interactome of *FLT3-ITD vs FLT3*-wildtype AML cells (Benjamini-Hochberg adjusted p-value <0.05) (heatmap, counts in Table S4, normalized to CEBPA amounts in the same cells). **B) CEBPA interactions with UHRF1 and USP7 were also observed by IP-Western blot (IP-WB); inhibiting FLT3-ITD signaling with sunitinib 50nM increased CEBPA interactions with USP7 and decreased interactions with UHRF1**. **C) Reverse IP-LCMS/MS analyses of USP7 in a different** *FLT3-ITD* **AML cell line (MV4-11).** IP-LCMS/MS of endogenous USP7 from *FLT3-ITD* AML cells (MV4-11), methods per panel A. Impact of inhibiting FLT3-ITD signaling with sunitinib 50nM on CEBPA and UHRF1 interactions was measured by total spectrum counts (Table S5) normalized to USP7 in the same sample. **D) Inhibiting FLT3-ITD signaling with sunitinib 50nM decreased USP7 phosphorylation at highly conserved serine-18 (S18-P)**. Mass spectrometry as per panel C (spectra in Figure S2). **E) Time-course decreases in USP7 S18-P and CEBPA S21-P measured by Western blots, with concurrent increases in total CEBPA, USP7 and p53 protein, in MOLM13 cells treated with sunitinib 50nM.** Since USP7 is known to regulate p53 in other cells, we also measured p53 protein. Whole cell lysates. Bar graphs show densitometric analysis performed using ImageJ software and normalized to actin in the same sample.

To corroborate the CEBPA interactions with UHRF1 and USP7 detected by mass-spectrometry, we IPed endogenous CEBPA from *FLT3-ITD* MOLM13 AML cells and performed Western blots (IP-WB), confirming strong signals for both UHRF1 and USP7 (**Figure 2B**). To examine impacts of FLT3-ITD signaling on these interactions, we treated the cells with the TKI sunitinib (50nM): TKI treatment decreased UHRF1, and increased USP7, pulled down with CEBPA (**Figure 2B**).

To further corroborate this data, we performed a reverse IP-LCMS/MS analyses of endogenous USP7 in a second *FLT3-ITD* AML cell line, MV4-11. As expected, major proteosome components, including UHRF1, were highly represented in the USP7 interactome (**Figure 2C**, **Table S5**). Inhibiting FLT3-ITD signaling in the cells with the TKI sunitinib 50nM introduced CEBPA into the USP7 interactome, and decreased the amount of UHRF1 (**Figure 2C, Table S5**). The LCMS/MS analyses also identified a post-translational phosphorylation modification of USP7 at serine-18 (S18-P), a residue highly conserved in evolution (**Figure 2D, S2**). Inhibiting FLT3-ITD signaling with the sunitinib decreased S18-P by ∼50% (**Figure 2D**). We used Western blots to corroborate this data in samples harvested serially over a 1 to 72 hour time-span: treating *FLT3-ITD* MOLM13 cells with sunitinib 50nM produced time-course decreases in both USP7 S18-P and CEBPA S21-P (a post-translational modification of CEBPA that is under FLT3-ITD signaling control(21)) (**Figure 2E**), and concurrently increased the total proteins (**Figure 2E**). Since USP7 (previously known as HAUSP) is known to stabilize p53 (the MTF regulator of apoptosis) in other cell contexts (30–32), this suggested USP7 stabilization might in turn also stabilize p53(30–32): inhibiting FLT3-ITD signaling with sunitinib increased USP7 and p53 proteins in parallel (**Figure 2E**).

### Inhibiting FLT3-ITD signaling decreased USP7 S18-P and CEBPA S21-P and increased the total proteins

To extend this data further, we treated a different *FLT3-ITD* AML cell line, MV4-11, with the TKI sunitinib to inhibit FLT3-ITD signaling, and measured phosphorylated and total USP7 and CEBPA protein levels by Western blots serially over a 1 to 72 hour time-span. Inhibiting FLT3-ITD signaling in MV4-11 cells with sunitinib produced time-course decreases in both USP7 S18-P and CEBPA S21-P (**Figure 3A**) and concurrently increased the total proteins (**Figure 3A**).

**Figure 3.**
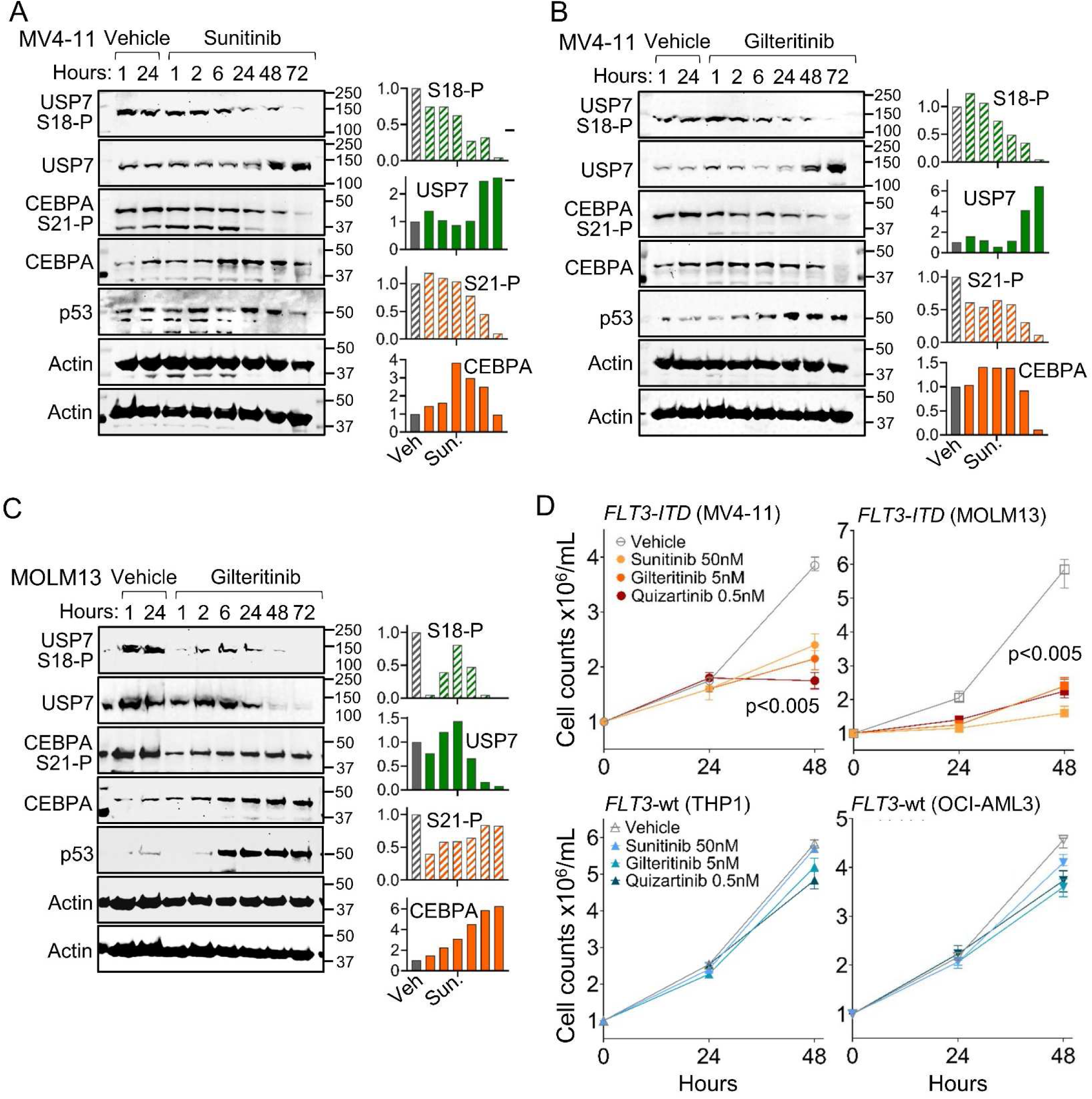
Inhibiting FLT3-ITD signaling with small molecule tyrosine kinase inhibitors (TKI) decreased USP7 S18-P and CEBPA S21-P, and concurrently increased total CEBPA, USP7 and p53 proteins. **A) Time-course decrease in USP7 S18-P and CEBPA S21-P, with concurrent increases in total CEBPA and USP7 protein, in *FLT3-ITD* MV4-11 cells treated with sunitinib 50nM.** Western blots, whole cell lysates. Bar graphs show densitometric analysis performed using ImageJ software and normalized to actin in the same sample. **B) Extension of this data to** *FLT3-ITD* **AML cells MV4-11 treated with a different TKI, gilteritinib 20nM**. **C) Extension of this data to** *FLT3-ITD* **AML cells MOLM13 treated with gilteritinib 20nM**. **D) Several different TKI significantly decreased proliferation of** *FLT3-ITD* **(MV4-11, MOLM13) but not** *FLT3*-wildtype **(THP1, OCI-AML3) AML cells.** Cell counts by automated counter, mean±standard deviation for 3 independent replicate experiments, relative to baseline cell count. p-values paired t-test, 2-sided, for cell counts at the 48 hour time-point, significant only in the *FLT3-ITD* AML cells.

We further extended this data by using a different TKI, gilteritinib, to inhibit FLT3-ITD signaling in the two *FLT3-ITD* AML cell lines MV4-11 and MOLM13: gilteritinib decreased USP7 S18-P and CEBPA S21-P (**Figure 3B, C**), and concurrently increased the total proteins (**Figure 3C, D**), in both *FLT3-ITD* AML cell lines. By contrast, TKI treatment of *FLT3*-wildtype AML cells (THP1) neither decreased CEBPA S21-P nor increased total CEBPA (**Figure S3A, B**).

In both *FLT3-ITD* AML cell lines, both TKIs also increased USP7 and p53 protein in parallel (**Figure 3A-C**). These data predicted genetic loss-of-function to *TP53* is redundant in the *FLT3-ITD* context, verified by mutual exclusivity of *TP53*- and *FLT3-ITD* mutations in clinical series (**Figure S4**).

Since CEBPA and p53 normally drive to terminal-differentiation and apoptosis these data suggested that the TKIs should have anti-proliferative effects in *FLT3-ITD* AML cells in particular. Accordingly, three different clinical TKIs to inhibit FLT3-ITD signaling, sunitinib, gilteritinib or quizartinib, all significantly decreased the proliferation of *FLT3-ITD* AML cells MV4-11 and MOLM13 (**Figure 3D**) but not *FLT3*-wildtype AML cells THP1 or OCI-AML3 (**Figure 3D**).

### Parsing cell fate consequences

Since CEBPA and p53 drive to cell fates of lineage-differentiation and apoptosis respectively, we measured for these specific cell fates in *FLT3-ITD* AML cells (MOLM13) with *FLT3*-wildtype (THP1) AML cells as a control. Sunitinib to inhibit FLT3-ITD signaling increased CEBPA nuclear protein levels in the *FLT3-ITD* cells, measured by immunofluorescence microscopy to complement the earlier Western blot data (**Figure 4A**). By contrast, CEBPA protein was abundant at baseline in the *FLT3*-wildtype AML cells and was not increased further by sunitinib treatment (**Figure 4B**). As expected from an increase in CEBPA in the *FLT3-ITD* (MOLM13) but not *FLT3*-wildtype (THP1) AML cells, there were morphologic changes of granulocytic differentiation (nuclear condensation and segmentation, cytoplasmic granulation and vacuolization, decrease in nuclear:cytoplasm ratio) in the *FLT3-ITD* (**Figure 4C**) but not *FLT3*-wildtype AML cells (**Figure 4D**). Corroborating the morphology data, a dose-dependent increase in the granulo-monocytic lineage-differentiation marker CD11b was observed in sunitinib-treated MOLM13 cells, measured by flow cytometry (**Figure 4E**). Apoptosis was also activated measured by flow cytometry from Annexin-V and propidium iodide (PI) staining (**Figure 4F**). Thus, the TKI-induced decrease in proliferation of *FLT3-ITD* but not *FLT3*-wildtype AML cells (**Figure 4G**) was via both resumed lineage-differentiation and apoptosis.

**Figure 4.**
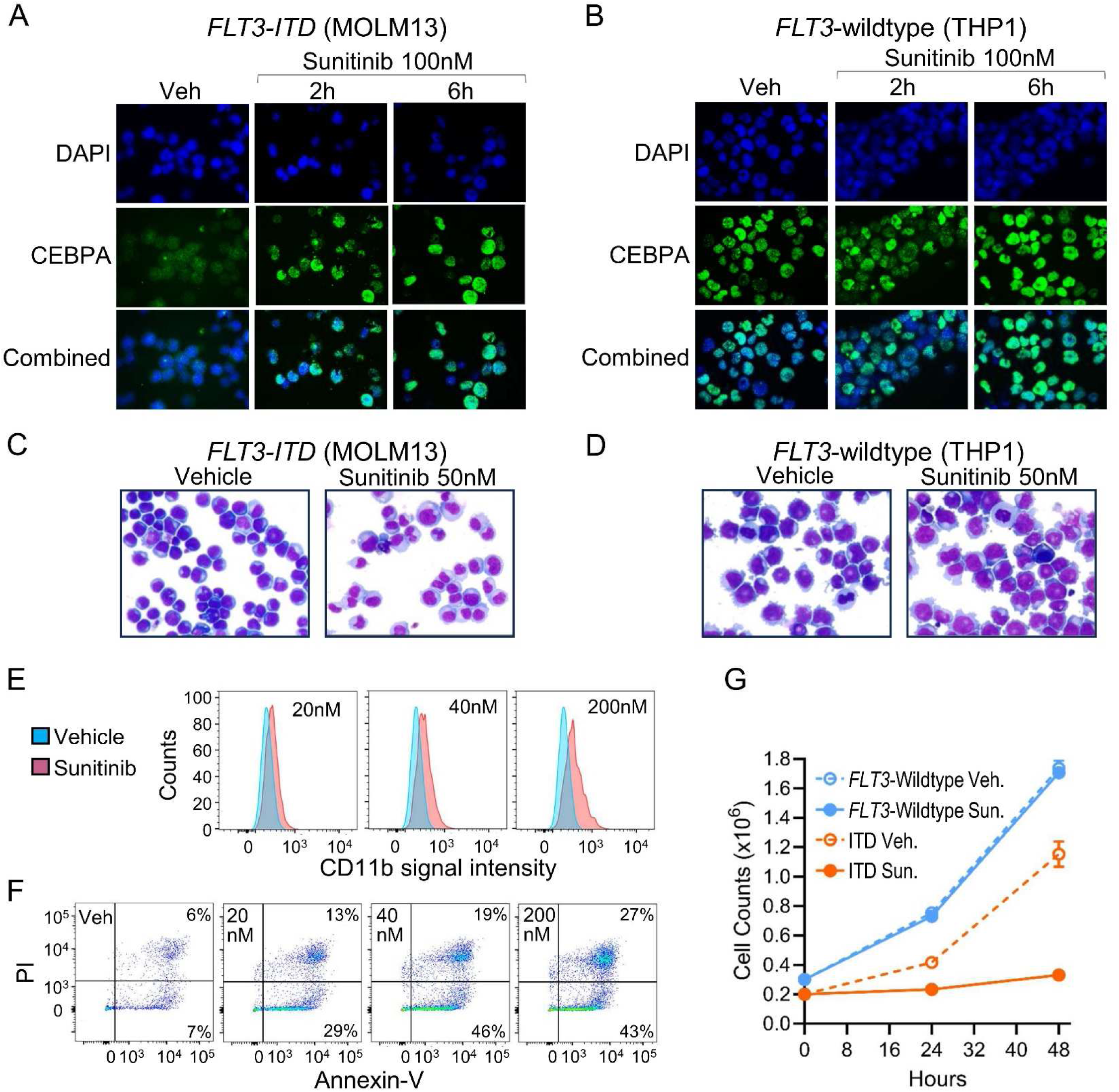
Inhibiting FLT3-ITD signaling restored CEBPA protein and activated lineage-differentiation and apoptosis in *FLT3-ITD* but not *FLT3*-wildtype (wt) AML cells. **A) Inhibiting FLT3-ITD signaling with sunitinib 100nM increased CEBPA protein in *FLT3-ITD* (MOLM13) AML cells.** Nuclei were stained with DAPI. Images by Nikon Eclipse 400 microscope; original magnification, ×630. **B) CEBPA protein was abundant at baseline in** *FLT3*-wt **AML cells (THP1) and was not further increased by sunitinib 100nM. C, D) Sunitinib treatment produced morphologic changes of granulocyte lineage-differentiation in** *FLT3-ITD* **(panel C) but not** *FLT3*-wildtype **AML cells (panel D)** (nuclear segmentation, increase in cytoplasmic neutrophilic granules, decrease in nuclear:cytoplasm ratio). Methods as per panel A. Giemsa-stained cytospin preparations after 48 hours of sunitinib treatment. Leica DMR microscope, magnification 400X. **E) The granulo-monocytic lineage-differentiation marker CD11b increased in the** *FLT3-ITD* **AML cells.** A sunitinib dose-effect was observed. Flow cytometry 48 hours after sunitinib addition. **F) Apoptosis was also activated in a dose- dependent manner in** *FLT3-ITD* **AML cells.** Annexin-V and propidium iodide (PI) staining measured by flow cytometry. **G)** *FLT3-ITD* **(MOLM13), but not** *FLT3*-wildtype **(THP1), AML cell proliferation was terminated.** Cell counts by automated counter.

### Small molecule inhibitors of the UPP also increased CEBPA and p53 protein in *FLT3-ITD* AML cells

These pathway data implied that UPP-inhibitors should also restore the MTFs, but by contrast to TKI, also of the phosphorylated CEBPA form. We therefore treated the two *FLT3-ITD* AML cell lines, MV4-11 and MOLM13, with two different proteosome inhibitors, the clinical inhibitor of the chymotrypsin-like subunit of the 26S proteasome bortezomib, or the peptidyl-aldehyde inhibitor of the 20S proteosome MG-132, and measured the phosphorylated and total protein levels by Western blots serially over a 1 to 72 hour time-span.

Bortezomib or MG-132 added to MV4-11 cells increased total USP7, CEBPA, and p53 protein within 1 hour and peaking at ∼6 hours (**Figure 5A, B**). Contrasting with the TKI-treatment results, wherein USP7 S18-P and CEBPA S21-P decreased, UPP-inhibitor treatment increased USP7 S18-P and CEBPA S21-P (**Figure 5A, B**).

**Figure 5.**
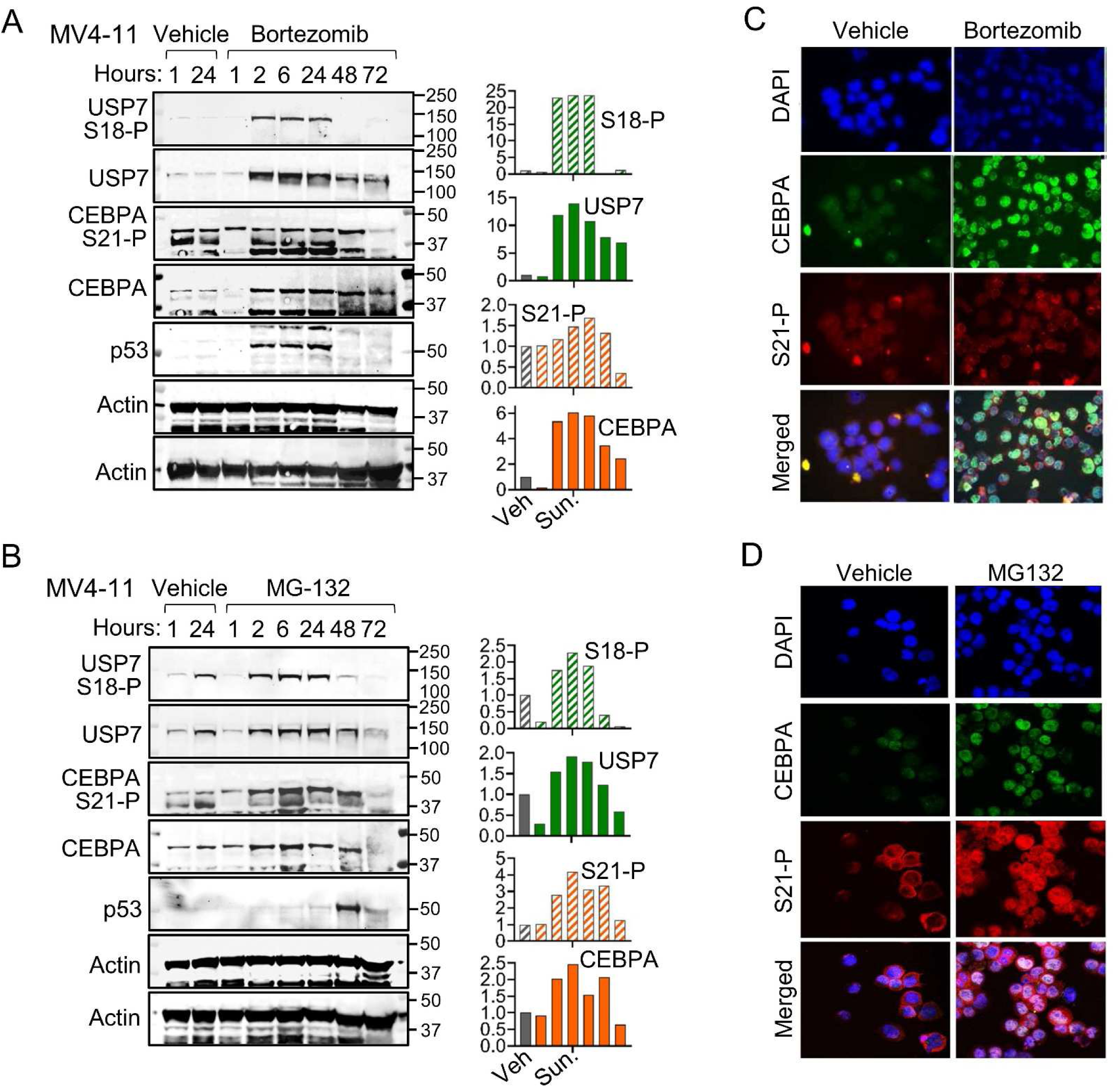
Small molecule inhibitors of the UPP increased CEBPA and p53 protein in *FLT3-ITD* AML cells MV4-11. **A, B) UPP-inhibitors bortezomib 5nM or MG132 1µM increased phosphorylated and total CEBPA, phosphorylated and total USP7, and p53 proteins in MV4-11 AML cells.** Western blots. There was extensive apoptosis at time-points after 24 hours. Bar graphs show densitometric analysis performed using ImageJ software and normalized to actin in the same sample. **C, D) Phosphorylated CEBPA increased in cytoplasm, whereas the total CEBPA increases were localized to nuclei**. *Treatments were* bortezomib 5nM (panel C) or MG-132 1µM (panel D) for 1 hour. Nuclei stained with DAPI. Images by Nikon Eclipse 400 microscope; original magnification, ×630.

We also examined the cellular location of CEBPA S21-P by immunofluorescence microscopy: the UPP-inhibitor treatments increased CEBPA S21-P in cytoplasm, contrasting with nuclear location of total CEBPA (**Figure 5C, D**).

The same treatments were applied to a second *FLT3-ITD* AML cell line, MOLM13 (**Figure 6A, B**): bortezomib or MG-132 treatments increased phosphorylated- and total-CEBPA and USP7, and total p53, in MOLM13 cells.

**Figure 6.**
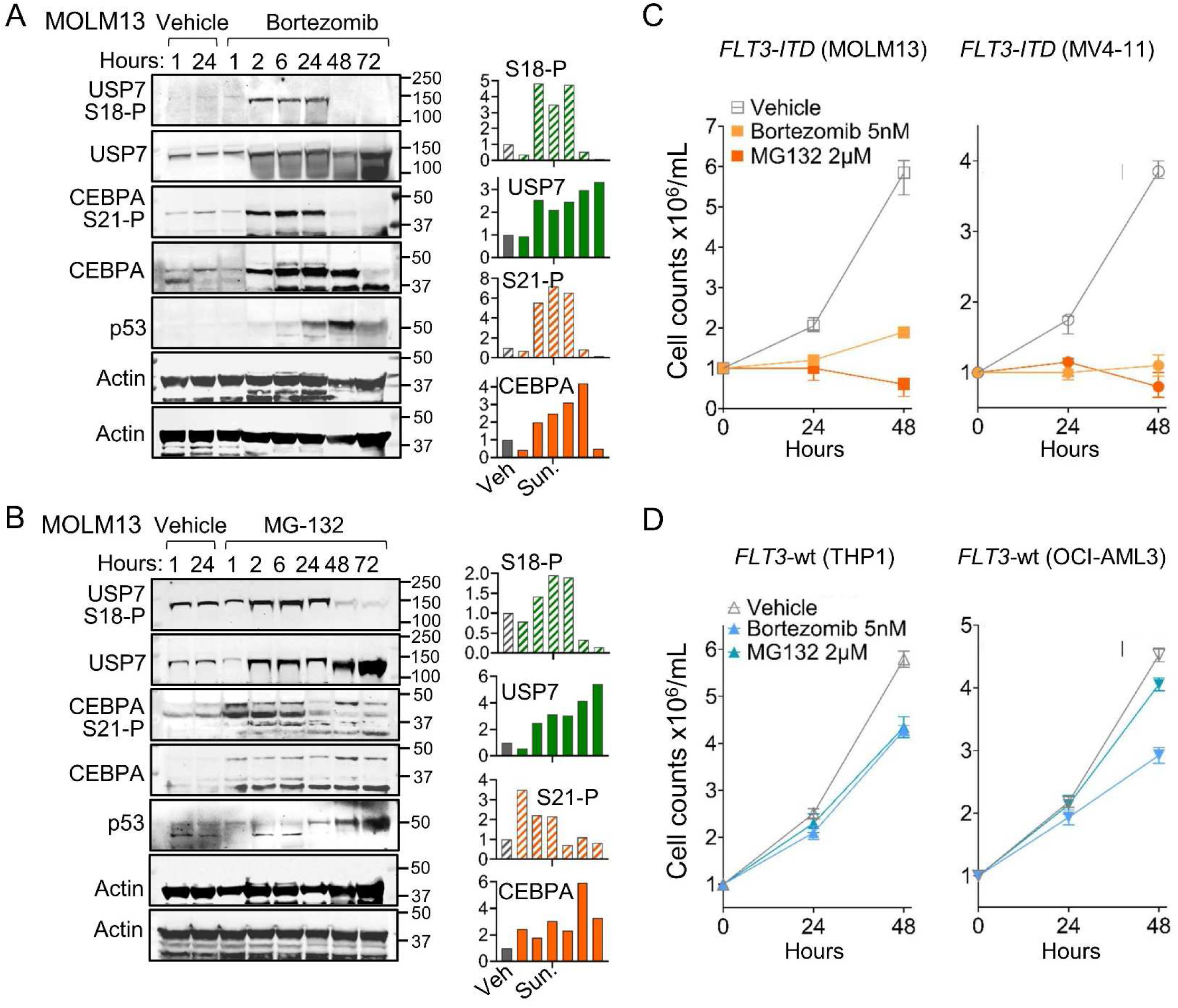
**Small molecule inhibitors of the UPP increased CEBPA and p53 protein in a second *FLT3-ITD* AML cell line MOLM13 also, and *FLT3-ITD* AML cells MV4-11 and MOLM13 were more sensitive to UPP-inhibitors than FLT3-wildtype AML cells THP1 and OCI-AML3**. **A, B) UPP-inhibitors bortezomib 5 nM or MG132 1 µM increased phosphorylated and total CEBPA, phosphorylated and total USP7, and p53 proteins in** *FLT3-ITD* **AML cells (MOLM13).** Western blots. There was extensive apoptosis at time-points after 24 hours. Bar graphs shows densitometric analysis performed using ImageJ software and normalized to actin in the same sample. **C, D)** *FLT3-ITD* **AML cells (MV4-11, MOLM13) were more sensitive to growth inhibition by UPP-inhibitors than** *FLT3*-wt **AML cells (THP1, OCI-AML3).** UPP-inhibitors were bortezomib 5nM and MG132 2µM. Cell counts by automated counter, mean ± standard deviation for 3 independent replicate experiments, relative to baseline cell count.

Bortezomib or MG-132 treatments of *FLT3*-wildtype AML cells (THP1, OCI-AML3) did not increase CEBPA or CEBPA S21-P protein (**Figure S5**).

As expected from the MTF protein rescue, bortezomib or MG-132 treatments decreased proliferation of *FLT3-ITD* AML cells (MV4-11, MOLM13) several-fold more than *FLT3*-wildtype AML cells (THP1, OCI-AML3)(**Figure 6C, D**).

### UPP-inhibitors bortezomib or MG132 terminated proliferation of gilteritinib-resistant *FLT3-ITD* AML cells

TKI-resistance is a clinical problem for patients with *FLT3-ITD* AMLs. Gilteritinib-resistant *FLT3-ITD* AML cells (MOLM13) were selected by continuous culture in gilteritinib 10nM (**Figure 7A**). Adding the UPP-inhibitors bortezomib at a clinically relevant concentration of 20nM (clinical plasm C_max_ is ∼300nM), or MG-132 0.5µM upregulated CEBPA and p53-proteins measured by Western blots – the CEBPA protein increase with UPP-inhibitors versus vehicle control was not as marked as seen in parental MOLM13 cells because of upregulation already present in the vehicle control by the continuous culture in gilteritinib (**Figure 7B**). The treated cells upregulated the monocytic lineage-differentiation markers CD14 and CD86, measured by flow cytometry – that is, the lineage-differentiation was in a monocytic direction contrasting with the granulo-monocytic direction seen in parental MOLM13 cells treated with UPP-inhibitors (**Figure 7C**). Morphologic changes of monocytic lineage-differentiation - nuclear condensation, increase in cell size, decrease in nuclear:cytoplasm ratio – were also observed in Giemsa-stained cytospin preparations (**Figure 7D**). The treatments also activated apoptosis measured by caspase 3/7 activation assays (**Figure 7E**). As expected from these effects, proliferation of the gilteritinib-resistant MOLM13 cells was significantly decreased by the UPP-inhibitor treatments (**Figure 7F**).

**Figure 7.**
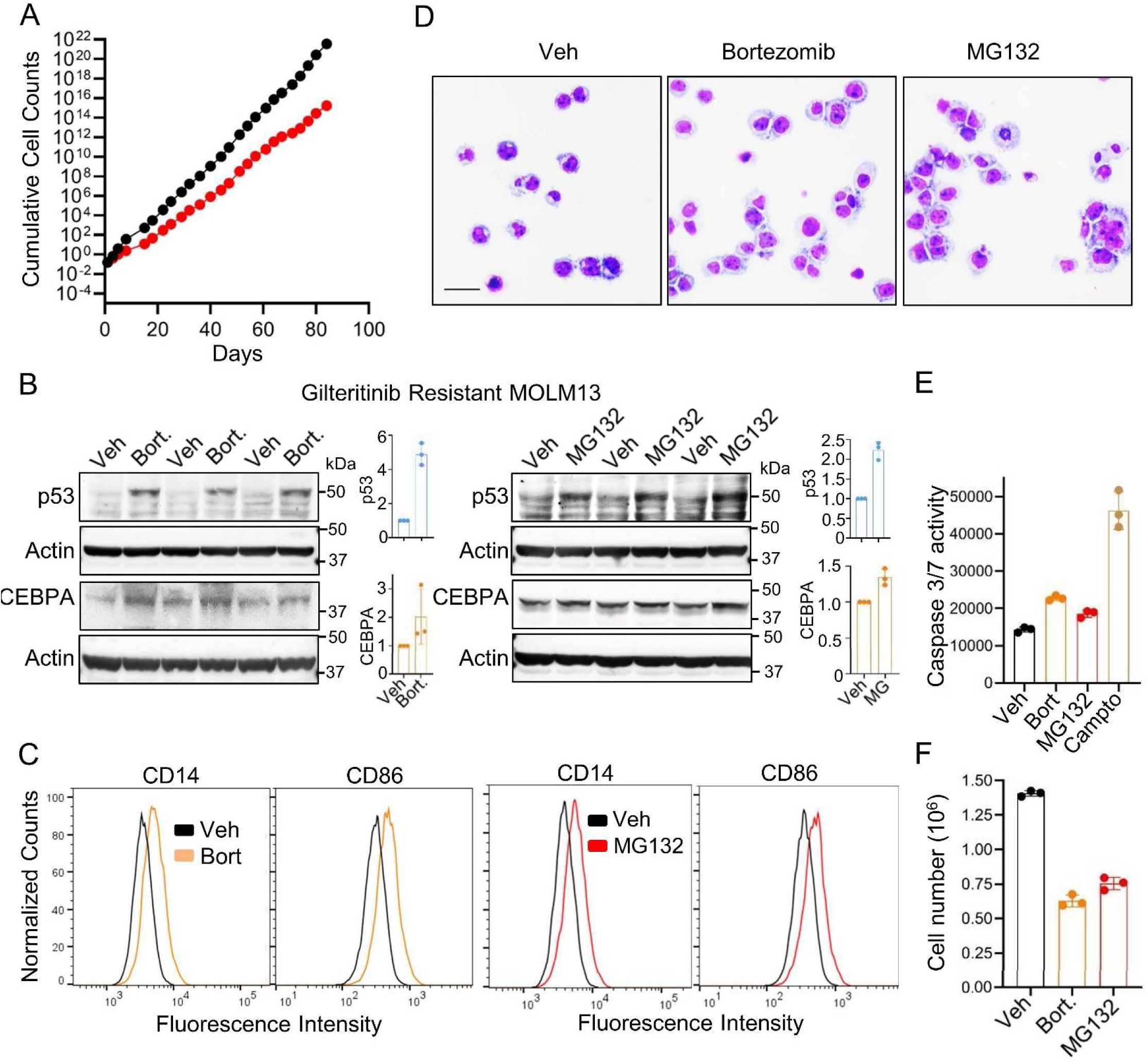
UPP-inhibitor treatment of TKI-resistant *FLT3-ITD* AML cells. **A) Gilteritinib-resistant *FLT3-ITD* AML cells (MOLM13) were selected by continuous culture in gilteritinib 10nM**. Media containing gilteritinib was replaced every 72 hours, with cell counts by automated counter each time. **B) UPP-inhibitors bortezomib 20nM (clinically relevant concentration) or MG-132 0.5µM upregulated CEBPA and p53-proteins in TKI-resistant MOLM13 cells.** CEBPA was already relatively high in the vehicle control TKI-resistant MOLM13 because of the continuous culture in gilteritinib. Western blots. Bar graphs shows densitometric analysis performed using ImageJ software and normalized to actin in the same sample. Experiments in triplicate. **C) The monocytic lineage-differentiation markers CD14 and CD86 increased, measured by flow cytometry 72 hours after addition of bortezomib 20nM** (left panels) **or MG132 0.5µM** (right panels). The continuous culture in gilteritinib had already upregulated the granulocytic marker CD11b. **D) Morphologic changes of monocytic lineage-differentiation - nuclear condensation, increase in cell size, decrease in nuclear:cytoplasm ratio – were observed in Giemsa-stained cytospin preparations**. Cytospins 72 hours after bortezomib or MG132 addition. Leica DMR microscope, magnification 400X, scale bar 20 μm. **E) Apoptosis was also activated measured by caspase 3/7 activation assay.** Measured 24 hours after UPP-inhibitor addition. **F) The UPP-inhibitor treatments significantly decreased proliferation of the gilteritinib-resistant MOLM13 cells**. Cell counts by automated counter.

## DISCUSSION

Experimental knockdown of *CEBPA* in GMPs decouples lineage-maturation from proliferation to expand myeloblasts (20, 48), and mutations that truncate and/or prevent DNA-binding by CEBPA are highly recurrent in human AMLs (45). Stated simply, loss-of-function to CEBPA is leukemogenic, as is loss-of-function, or dominant-negative, impacts on the other lineage MTFs, RUNX1 or SPI1, with which CEBPA cooperates to drive granulo-monocytic lineage-differentiation (reviewed in (49)). Here we found that besides mutating *CEBPA*, leukemogenesis selects for another method to produce CEBPA loss-of-function – FLT3-ITD signaling that destabilizes CEBPA protein. We found high CEBPA mRNA but little CEBPA protein in *FLT3-ITD* versus *FLT3*-wildtype AML cells. Unbiased analyses of the CEBPA protein-protein interactome disclosed prominent interactions with major UPP components UHRF1 (a ubiquitin-ligase) and USP7 (a deubiquitinase), that were especially enriched in *FLT3-ITD* vs *FLT3*-wildtype AML cells. Inhibiting FLT3-ITD signaling decreased CEBPA and USP7 serine-phosphorylations, S21-P and S18-P respectively, shifted CEBPA interactions from UHRF1 to USP7 that is expected to stabilize CEBPA, and accordingly rescued CEBPA protein in the cells. CEBPA protein regulation by the UPP has been observed by others, finding signaling inputs from a tribbles family serine-threonine kinase TRIB2, and cyclin dependent kinase 2 (CDK2) (50, 51) - here we found FLT3-ITD signaling inputs into this system also.

Predicted from this data was that UPP-inhibitors, like TKIs to inhibit FLT3-ITD signaling, should increase total CEBPA but also CEBPA S21-P. UPP-inhibitors accordingly increased total CEBPA and CEBPA S21-P, with CEBPA S21-P in the cytoplasm and CEBPA in nuclei of *FLT3-ITD* AML cells. UPP-inhibitors produced these consequences even in TKI-resistant *FLT3-ITD* AML cells. Thus, a treatment implication of our observations is that UPP-inhibitors, which are presently used to treat other cancers but not AMLs (34, 35), could be candidate rational therapeutics to remedy the FLT3-ITD transforming pathway at points further downstream than engaged by TKI. Supporting this proposition, others have also observed specific sensitivity of *FLT3-ITD* vs *FLT3*-wildtype AML cells to UPP-inhibitors (34, 35, 51).

TKIs to inhibit FLT3-ITD signaling, when combined with another standard AML treatment modality of the HMAs, has been accompanied by clinical ‘differentiation syndrome’ (52–54) - synchronous differentiation of *FLT3-ITD* myeloblasts into neutrophils resulting in inflammatory symptoms and signs such as neutrophilic dermatoses (22, 55, 56). Such differentiation requires and is driven by CEBPA. Therefore, the present mechanism data suggests an underlying mechanism for therapeutic cooperation between HMAs, which inhibit a repressing epigenetic enzyme DNMT1 to permit AML cells to activate onward lineage-differentiation programs (57–60), and TKIs (reviewed in (61)).

These links between FLT3-ITD and CEBPA also explain mutual exclusivity of *FLT3-ITD* and *CEBPA*^dm^ in clinical AML series, and absence of *FLT3-ITD* in AMLs with erythro-megakaryocytic phenotypes, lineages in which CEBPA is not normally expressed (12).

*TP53* is the single most frequently mutated and deleted gene in cancers. This is a major problem in oncology because chemotherapy treatments aim to upregulate its protein product p53 to activate apoptosis. *TP53* is normally continuously transcribed in proliferating cells, e.g., myeloid progenitors, and p53 protein levels are instead kept in check by continuous proteolysis, promoted by the ubiquitin ligases MDM2/MDM4 (25–28) while USP7, a deubiquitinase, participates in stabilizing p53 (29–32). *MDM2* or *MDM4* amplifications are thus another method besides *TP53* mutations by which cancers suppress p53 function. Here we found yet another UPP-centered method for suppressing p53 function in malignant transformations: FLT3-ITD signaling destabilized USP7 and p53: inhibiting *FLT3-ITD* signaling stabilized USP7 and p53 proteins, and activated apoptosis as well as CEBPA-driven granulo-monocytic differentiation in *FLT3-ITD* but not *FLT3*-wildtype AML cells. This connection between FLT3-ITD signaling and p53 status explains: (i) mutual exclusivity of *FLT3-ITD* and *TP53*-mutations (62); (ii) clinical chemoresistance of *FLT3-ITD* AMLs - even with chemotherapy intensification above usual levels, *FLT3-ITD* AMLs relapse early and at high-rates (8, 63–65); FLT3-ITD signaling impacts on CEBPA function is an unlikely mechanism for chemo-resistance, because *CEBPA*^dm^ AMLs, which also have loss of CEBPA transactivating functions, are chemo-sensitive (45); (iii) synergy between chemotherapy and TKIs both in vitro and in the clinic (52, 53, 66); and (iv) UPP-inhibitor activity against *FLT3-ITD* AML cells in particular compared to *FLT3*-wildtype AML cells, observed in vitro here by us and also seen in vitro previously by others (34, 35, 51).

Other class III tyrosine kinase receptors, KIT, PDGFRA, PDGFRB, CSF1R, also expand specific myeloid lineage compartments when ligand-activated. The genes for these receptors are also recurrently mutated and/or translocated in myeloid malignancies, accompanied by distinctive lineage-features of disease (67). Conceivably, the oncoprotein forms of these other class III tyrosine kinase receptors also signal into the UPP to destabilize MTFs corresponding to the specific lineage-contexts and/or p53. Supporting these possibilities, USP7 was shown to stabilize the myeloid lineage MTF GATA1 that is essential for erythroid differentiation (68), and these other myeloid malignancies are also typically chemo-refractory.

### Limitations

A major limitation is whether results observed in AML cell lines, model systems widely used for practical reasons, apply to AML cells in patients. To support extrapolation/generalizability, corroborating and matching data was generated by TKI treatment of primary AML cells and by phenotype, gene expression and DNA-sequencing analyses in large clinical AML sample cohorts. Also supporting validity and generalizability of the identified core mechanisms they explain and fit with multiple facets of *FLT3-ITD* AMLs, including GMP cell-of-origin, GMP phenotype of myeloblasts, experimental leukemogenesis by CEBPA knock-down in GMPs, lineage/maturation features of primary *FLT3-ITD* myeloblasts, clinical chemotherapy-resistance of *FLT3-ITD* AMLs in the absence of *TP53*-mutations, neutrophilic differentiation syndrome with clinical TKI + HMA therapy, and specific sensitivity of *FLT3-ITD* AML cells to UPP-inhibitors.

In sum, FLT3-ITD signaling promotes UPP-mediated proteolysis of CEBPA and p53, MTFs that normally terminate GMP replications via lineage-maturation and apoptosis respectively. Candidate remedies for this leukemogenic pathway, other than TKI, are UPP-inhibitors of which several are approved to treat other malignancies.

## AUTHOR CONTRIBUTION STATEMENT

X.G., Z.Z., S.B., S.B., R.B., J.M., B.J., and Y.S. performed experiments and research. X.G., Z.Z., S.B., S.B., B.J., and Y.S. analyzed data. X.G. and Y.S. generated hypotheses and designed research. Y.S. obtained funding and wrote the paper. All authors reviewed/edited the manuscript.

## ACKNOWLEDGEMENTS

We acknowledge administrative support from JoAnn Bandera, and facility support from the Biological Resources Unit (BRU), FlowCore, and Imaging Core of the Cleveland Clinic Lerner Research Institute. YS is supported by National Heart, Lung and Blood Institute PO1 HL146372; National Cancer Institute P30 CA043703; RO1 CA204373; R21 CA263430, philanthropic funds from Robert and Jennifer McNeil, Leszek and Jolanta Czarnecki, and Dane and Louise Miller, and the James Oberle family, and NIH Shared Instrument award S10OD018205.

## REFERENCES

1. Nakao M, Yokota S, Iwai T, Kaneko H, Horiike S, Kashima K, et al. Internal tandem duplication of the flt3 gene found in acute myeloid leukemia. Leukemia. 1996;10(12):1911–8.

2. Kiyoi H, Towatari M, Yokota S, Hamaguchi M, Ohno R, Saito H, et al. Internal tandem duplication of the FLT3 gene is a novel modality of elongation mutation which causes constitutive activation of the product. Leukemia. 1998;12(9):1333–7.

3. Koch S, Jacobi A, Ryser M, Ehninger G, and Thiede C. Abnormal localization and accumulation of FLT3-ITD, a mutant receptor tyrosine kinase involved in leukemogenesis. Cells Tissues Organs. 2008;188(1-2):225–35.

4. Choudhary C, Olsen JV, Brandts C, Cox J, Reddy PN, Bohmer FD, et al. Mislocalized activation of oncogenic RTKs switches downstream signaling outcomes. Mol Cell. 2009;36(2):326–39.

5. Mizuki M, Fenski R, Halfter H, Matsumura I, Schmidt R, Muller C, et al. Flt3 mutations from patients with acute myeloid leukemia induce transformation of 32D cells mediated by the Ras and STAT5 pathways. Blood. 2000;96(12):3907–14.

6. Yamawaki K, Shiina I, Murata T, Tateyama S, Maekawa Y, Niwa M, et al. FLT3-ITD transduces autonomous growth signals during its biosynthetic trafficking in acute myelogenous leukemia cells. Sci Rep. 2021;11(1):22678.

7. Levis M, and Perl AE. Gilteritinib: potent targeting of FLT3 mutations in AML. Blood advances. 2020;4(6):1178–91.

8. Erba HP, Montesinos P, Kim HJ, Patkowska E, Vrhovac R, Zak P, et al. Quizartinib plus chemotherapy in newly diagnosed patients with FLT3-internal-tandem-duplication-positive acute myeloid leukaemia (QuANTUM-First): a randomised, double-blind, placebo-controlled, phase 3 trial. Lancet. 2023;401(10388):1571–83.

9. Jan M, Snyder TM, Corces-Zimmerman MR, Vyas P, Weissman IL, Quake SR, et al. Clonal evolution of preleukemic hematopoietic stem cells precedes human acute myeloid leukemia. Sci Transl Med. 2012;4(149):149ra18.

10. Corces-Zimmerman MR, Hong WJ, Weissman IL, Medeiros BC, and Majeti R. Preleukemic mutations in human acute myeloid leukemia affect epigenetic regulators and persist in remission. Proc Natl Acad Sci U S A. 2014;111(7):2548–53.

11. Shlush LI, Zandi S, Mitchell A, Chen WC, Brandwein JM, Gupta V, et al. Identification of pre-leukaemic haematopoietic stem cells in acute leukaemia. Nature. 2014;506(7488):328–33.

12. Zeng AGX, Iacobucci I, Shah S, Mitchell A, Wong G, Bansal S, et al. Single-cell Transcriptional Atlas of Human Hematopoiesis Reveals Genetic and Hierarchy-Based Determinants of Aberrant AML Differentiation. Blood Cancer Discov. 2025:OF1-OF18.

13. Bottomly D, Long N, Schultz AR, Kurtz SE, Tognon CE, Johnson K, et al. Integrative analysis of drug response and clinical outcome in acute myeloid leukemia. Cancer cell. 2022;40(8):850–64 e9.

14. Li Y, Yang W, Patel RM, Casey EB, Denby E, Mendoza-Castrejon J, et al. FLT3ITD drives context-specific changes in cell identity and variable interferon dependence during AML initiation. Blood. 2023;141(12):1442–56.

15. Kim T, Lee H, Capo-Chichi JM, Chang MH, Yoo YS, Basi G, et al. Single cell proteogenomic sequencing identifies a relapse-fated AML subclone carrying FLT3-ITD with CN-LOH at chr13q. EJHaem. 2022;3(2):426–33.

16. Horiike S, Yokota S, Nakao M, Iwai T, Sasai Y, Kaneko H, et al. Tandem duplications of the FLT3 receptor gene are associated with leukemic transformation of myelodysplasia. Leukemia. 1997;11(9):1442–6.

17. Daver N, Strati P, Jabbour E, Kadia T, Luthra R, Wang S, et al. FLT3 mutations in myelodysplastic syndrome and chronic myelomonocytic leukemia. Am J Hematol. 2013;88(1):56–9.

18. Largeaud L, Bertoli S, Berard E, Tavitian S, Picard M, Dufrechou S, et al. Genomic landscape of hyperleukocytic acute myeloid leukemia. Blood cancer journal. 2022;12(1):4.

19. Vierbuchen T, and Wernig M. Molecular roadblocks for cellular reprogramming. Mol Cell. 2012;47(6):827–38.

20. Theilgaard-Monch K, Pundhir S, Reckzeh K, Su J, Tapia M, Furtwangler B, et al. Transcription factor-driven coordination of cell cycle exit and lineage-specification in vivo during granulocytic differentiation : In memoriam Professor Niels Borregaard. Nat Commun. 2022;13(1):3595.

21. Radomska HS, Basseres DS, Zheng R, Zhang P, Dayaram T, Yamamoto Y, et al. Block of C/EBP alpha function by phosphorylation in acute myeloid leukemia with FLT3 activating mutations. J Exp Med. 2006;203(2):371–81.

22. Sexauer A, Perl A, Yang X, Borowitz M, Gocke C, Rajkhowa T, et al. Terminal myeloid differentiation in vivo is induced by FLT3 inhibition in FLT3/ITD AML. Blood. 2012;120(20):4205–14.

23. Mizuki M, Schwable J, Steur C, Choudhary C, Agrawal S, Sargin B, et al. Suppression of myeloid transcription factors and induction of STAT response genes by AML-specific Flt3 mutations. Blood. 2003;101(8):3164–73.

24. Zheng R, Friedman AD, Levis M, Li L, Weir EG, and Small D. Internal tandem duplication mutation of FLT3 blocks myeloid differentiation through suppression of C/EBPalpha expression. Blood. 2004;103(5):1883–90.

25. Prus G, Satpathy S, Weinert BT, Narita T, and Choudhary C. Global, site-resolved analysis of ubiquitylation occupancy and turnover rate reveals systems properties. Cell. 2024;187(11):2875–92 e21.

26. Abbas HA, Maccio DR, Coskun S, Jackson JG, Hazen AL, Sills TM, et al. Mdm2 is required for survival of hematopoietic stem cells/progenitors via dampening of ROS-induced p53 activity. Cell stem cell. 2010;7(5):606–17.

27. 27. Ringshausen I, O’Shea CC, Finch AJ, Swigart LB, and Evan GI. Mdm2 is critically and continuously required to suppress lethal p53 activity in vivo. Cancer cell. 2006;10(6):501–14.

28. Liu G, Terzian T, Xiong S, Van Pelt CS, Audiffred A, Box NF, et al. The p53-Mdm2 network in progenitor cell expansion during mouse postnatal development. The Journal of pathology. 2007;213(4):360–8.

29. Schauer NJ, Liu X, Magin RS, Doherty LM, Chan WC, Ficarro SB, et al. Selective USP7 inhibition elicits cancer cell killing through a p53-dependent mechanism. Sci Rep. 2020;10(1):5324.

30. Li M, Chen D, Shiloh A, Luo J, Nikolaev AY, Qin J, et al. Deubiquitination of p53 by HAUSP is an important pathway for p53 stabilization. Nature. 2002;416(6881):648–53.

31. Liu J, Cao L, Wang Y, Zou Y, Guo Q, Chen S, et al. The phosphorylation-deubiquitination positive feedback loop of the CHK2-USP7 axis stabilizes p53 under oxidative stress. Cell reports. 2024;43(6):114366.

32. Brooks CL, Li M, Hu M, Shi Y, and Gu W. The p53--Mdm2--HAUSP complex is involved in p53 stabilization by HAUSP. Oncogene. 2007;26(51):7262–6.

33. Konopleva M, Martinelli G, Daver N, Papayannidis C, Wei A, Higgins B, et al. MDM2 inhibition: an important step forward in cancer therapy. Leukemia. 2020;34(11):2858–74.

34. Larrue C, Saland E, Boutzen H, Vergez F, David M, Joffre C, et al. Proteasome inhibitors induce FLT3-ITD degradation through autophagy in AML cells. Blood. 2016;127(7):882–92.

35. Long J, Chen X, Shen Y, Lei Y, Mu L, Wang Z, et al. A combinatorial therapeutic approach to enhance FLT3-ITD AML treatment. Cell Rep Med. 2023;4(11):101286.

36. Gu X, Hu Z, Ebrahem Q, Crabb JS, Mahfouz RZ, Radivoyevitch T, et al. Runx1 regulation of Pu.1 corepressor/coactivator exchange identifies specific molecular targets for leukemia differentiation therapy. J Biol Chem. 2014;289(21):14881–95.

37. Gu X, Ebrahem Q, Mahfouz RZ, Hasipek M, Enane F, Radivoyevitch T, et al. Leukemogenic nucleophosmin mutation disrupts the transcription factor hub that regulates granulomonocytic fates. J Clin Invest. 2018;128(10):4260–79.

38. Hu Z, Gu X, Baraoidan K, Ibanez V, Sharma A, Kadkol S, et al. RUNX1 regulates corepressor interactions of PU.1. Blood. 2011;117(24):6498–508.

39. Galaxy C. The Galaxy platform for accessible, reproducible and collaborative biomedical analyses: 2022 update. Nucleic Acids Res. 2022;50(W1):W345–W51.

40. Lerdrup M, Johansen JV, Agrawal-Singh S, and Hansen K. An interactive environment for agile analysis and visualization of ChIP-sequencing data. Nat Struct Mol Biol. 2016;23(4):349–57.

41. Gislason MH, Demircan GS, Prachar M, Furtwangler B, Schwaller J, Schoof EM, et al. BloodSpot 3.0: a database of gene and protein expression data in normal and malignant haematopoiesis. Nucleic Acids Res. 2024;52(D1):D1138–D42.

42. Reich M, Liefeld T, Gould J, Lerner J, Tamayo P, and Mesirov JP. GenePattern 2.0. Nat Genet. 2006;38(5):500–1.

43. Tyner JW, Tognon CE, Bottomly D, Wilmot B, Kurtz SE, Savage SL, et al. Functional genomic landscape of acute myeloid leukaemia. Nature. 2018;562(7728):526–31.

44. Hasemann MS, Lauridsen FK, Waage J, Jakobsen JS, Frank AK, Schuster MB, et al. C/EBPalpha is required for long-term self-renewal and lineage priming of hematopoietic stem cells and for the maintenance of epigenetic configurations in multipotent progenitors. PLoS Genet. 2014;10(1):e1004079.

45. Georgi JA, Stasik S, Kramer M, Meggendorfer M, Rollig C, Haferlach T, et al. Prognostic impact of CEBPA mutational subgroups in adult AML. Leukemia. 2024;38(2):281–90.

46. Gu X, Enane F, Tohme R, Schuerger C, Radivoyevitch T, Parker Y, et al. PBRM1 loss in kidney cancer unbalances the proximal tubule master transcription factor hub to repress proximal tubule differentiation. Cell reports. 2021;36(12):109747.

47. Enane FO, Shuen WH, Gu X, Quteba E, Przychodzen B, Makishima H, et al. GATA4 loss of function in liver cancer impedes precursor to hepatocyte transition. J Clin Invest. 2017;127(9):3527–42.

48. Pundhir S, Bratt Lauridsen FK, Schuster MB, Jakobsen JS, Ge Y, Schoof EM, et al. Enhancer and Transcription Factor Dynamics during Myeloid Differentiation Reveal an Early Differentiation Block in Cebpa null Progenitors. Cell reports. 2018;23(9):2744–57.

49. Velcheti V, Schrump D, and Saunthararajah Y. Ultimate Precision: Targeting Cancer but Not Normal Self-replication. American Society of Clinical Oncology educational book American Society of Clinical Oncology Annual Meeting. 2018(38):950–63.

50. Thacker G, Mishra M, Sharma A, Singh AK, Sanyal S, and Trivedi AK. CDK2 destabilizes tumor suppressor C/EBPalpha expression through ubiquitin-mediated proteasome degradation in acute myeloid leukemia. Journal of cellular biochemistry. 2020;121(4):2839–50.

51. O’Connor C, Lohan F, Campos J, Ohlsson E, Salome M, Forde C, et al. The presence of C/EBPalpha and its degradation are both required for TRIB2-mediated leukaemia. Oncogene. 2016;35(40):5272–81.

52. Levis M, Pham R, Smith BD, and Small D. In vitro studies of a FLT3 inhibitor combined with chemotherapy: sequence of administration is important to achieve synergistic cytotoxic effects. Blood. 2004;104(4):1145–50.

53. Schittenhelm MM, Kampa KM, Yee KW, and Heinrich MC. The FLT3 inhibitor tandutinib (formerly MLN518) has sequence-independent synergistic effects with cytarabine and daunorubicin. Cell cycle. 2009;8(16):2621–30.

54. Ravandi F, Alattar ML, Grunwald MR, Rudek MA, Rajkhowa T, Richie MA, et al. Phase II study of azacytidine plus sorafenib in patients with acute myeloid leukemia and FLT-3 internal tandem duplication mutation. Blood. 2013.

55. McMahon CM, Canaani J, Rea B, Sargent RL, Qualtieri JN, Watt CD, et al. Gilteritinib induces differentiation in relapsed and refractory FLT3-mutated acute myeloid leukemia. Blood advances. 2019;3(10):1581–5.

56. Fathi AT, Le L, Hasserjian RP, Sadrzadeh H, Levis M, and Chen YB. FLT3 inhibitor-induced neutrophilic dermatosis. Blood. 2013;122(2):239–42.

57. Pinto A, Attadia V, Fusco A, Ferrara F, Spada OA, and Di Fiore PP. 5-Aza-2’-deoxycytidine induces terminal differentiation of leukemic blasts from patients with acute myeloid leukemias. Blood. 1984;64(4):922–9.

58. Negrotto S, Ng KP, Jankowska AM, Bodo J, Gopalan B, Guinta K, et al. CpG methylation patterns and decitabine treatment response in acute myeloid leukemia cells and normal hematopoietic precursors. Leukemia. 2012;26(2):244–54.

59. Ng KP, Ebrahem Q, Negrotto S, Mahfouz RZ, Link KA, Hu Z, et al. p53 independent epigenetic-differentiation treatment in xenotransplant models of acute myeloid leukemia. Leukemia. 2011;25(11):1739–50.

60. Saunthararajah Y, Sekeres M, Advani A, Mahfouz R, Durkin L, Radivoyevitch T, et al. Evaluation of noncytotoxic DNMT1-depleting therapy in patients with myelodysplastic syndromes. J Clin Invest. 2015;125(3):1043–55.

61. Zavras PD, Shastri A, Goldfinger M, Verma AK, and Saunthararajah Y. Clinical Trials Assessing Hypomethylating Agents Combined with Other Therapies: Causes for Failure and Potential Solutions. Clin Cancer Res. 2021;27(24):6653–61.

62. Sargas C, Ayala R, Larrayoz MJ, Chillon MC, Carrillo-Cruz E, Bilbao-Sieyro C, et al. Molecular Landscape and Validation of New Genomic Classification in 2668 Adult AML Patients: Real Life Data from the PETHEMA Registry. Cancers. 2023;15(2).

63. Bertoli S, Dumas PY, Berard E, Largeaud L, Bidet A, Delabesse E, et al. Outcome of Relapsed or Refractory FLT3-Mutated Acute Myeloid Leukemia Before Second-Generation FLT3 Tyrosine Kinase Inhibitors: A Toulouse-Bordeaux DATAML Registry Study. Cancers. 2020;12(4).

64. Burnett AK, Russell NH, Hills RK, and United Kingdom National Cancer Research Institute Acute Myeloid Leukemia Study G. Higher daunorubicin exposure benefits FLT3 mutated acute myeloid leukemia. Blood. 2016;128(3):449–52.

65. Meshinchi S, and Appelbaum FR. Structural and functional alterations of FLT3 in acute myeloid leukemia. Clin Cancer Res. 2009;15(13):4263–9.

66. Yee KW, Schittenhelm M, O’Farrell AM, Town AR, McGreevey L, Bainbridge T, et al. Synergistic effect of SU11248 with cytarabine or daunorubicin on FLT3 ITD-positive leukemic cells. Blood. 2004;104(13):4202–9.

67. Berenstein R. Class III Receptor Tyrosine Kinases in Acute Leukemia - Biological Functions and Modern Laboratory Analysis. Biomark Insights. 2015;10(Suppl 3):1–14.

68. Liang L, Peng Y, Zhang J, Zhang Y, Roy M, Han X, et al. Deubiquitylase USP7 regulates human terminal erythroid differentiation by stabilizing GATA1. Haematologica. 2019;104(11):2178–87.

69. Quentmeier H, Reinhardt J, Zaborski M, and Drexler HG. FLT3 mutations in acute myeloid leukemia cell lines. Leukemia. 2003;17(1):120–4.

70. Ghandi M, Huang FW, Jane-Valbuena J, Kryukov GV, Lo CC, McDonald ER, 3rd, et al. Next-generation characterization of the Cancer Cell Line Encyclopedia. Nature. 2019;569(7757):503–8.

