## Supplementary Figures for "FLT3-ITD signals for CEBPA and p53 proteolysis by the ubiquitin-proteosome pathway"

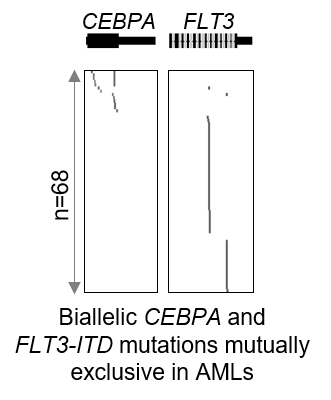

**Figure S1. *FLT3* mutations and *CEBPA* bi-allelic mutations are mutually exclusive.** Mutation calls from TCGA indicated by grey dots, primary AMLs with either *CEBPA* or *FLT3* mutations shown, n=68 (TCGA public data).

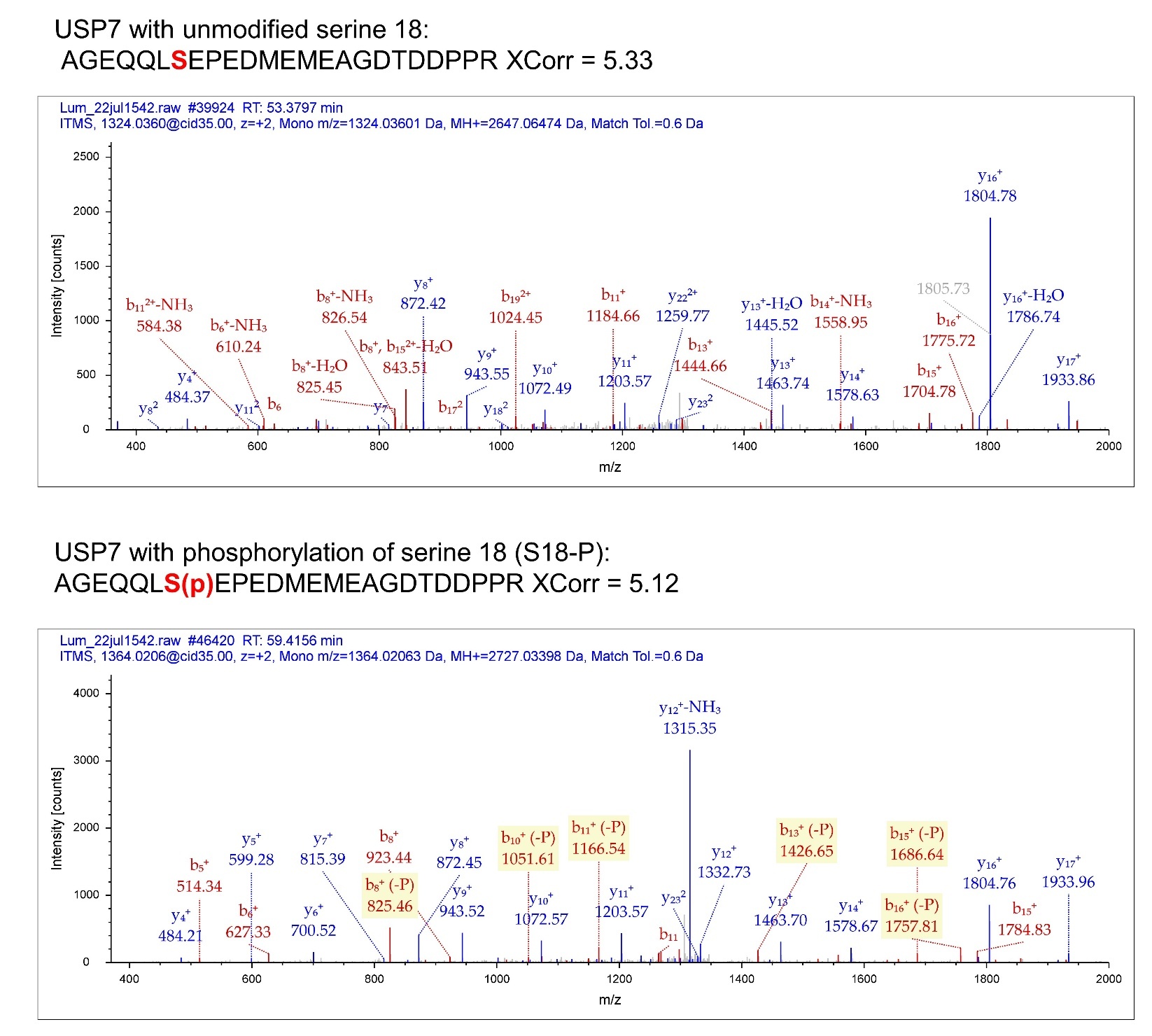

**Figure S2. USP7 phosphorylation at serine-18.** Tandem mass spectrum of unmodified and modified peptide AGEQQLSEPEDMEMEAGDTDDPPR. Phosphorylation at the position 7 serine in the peptide corresponds to serine 18 (S18) in USP7 protein. Xcorr = Sequest score that matches spectra to peptide sequences.

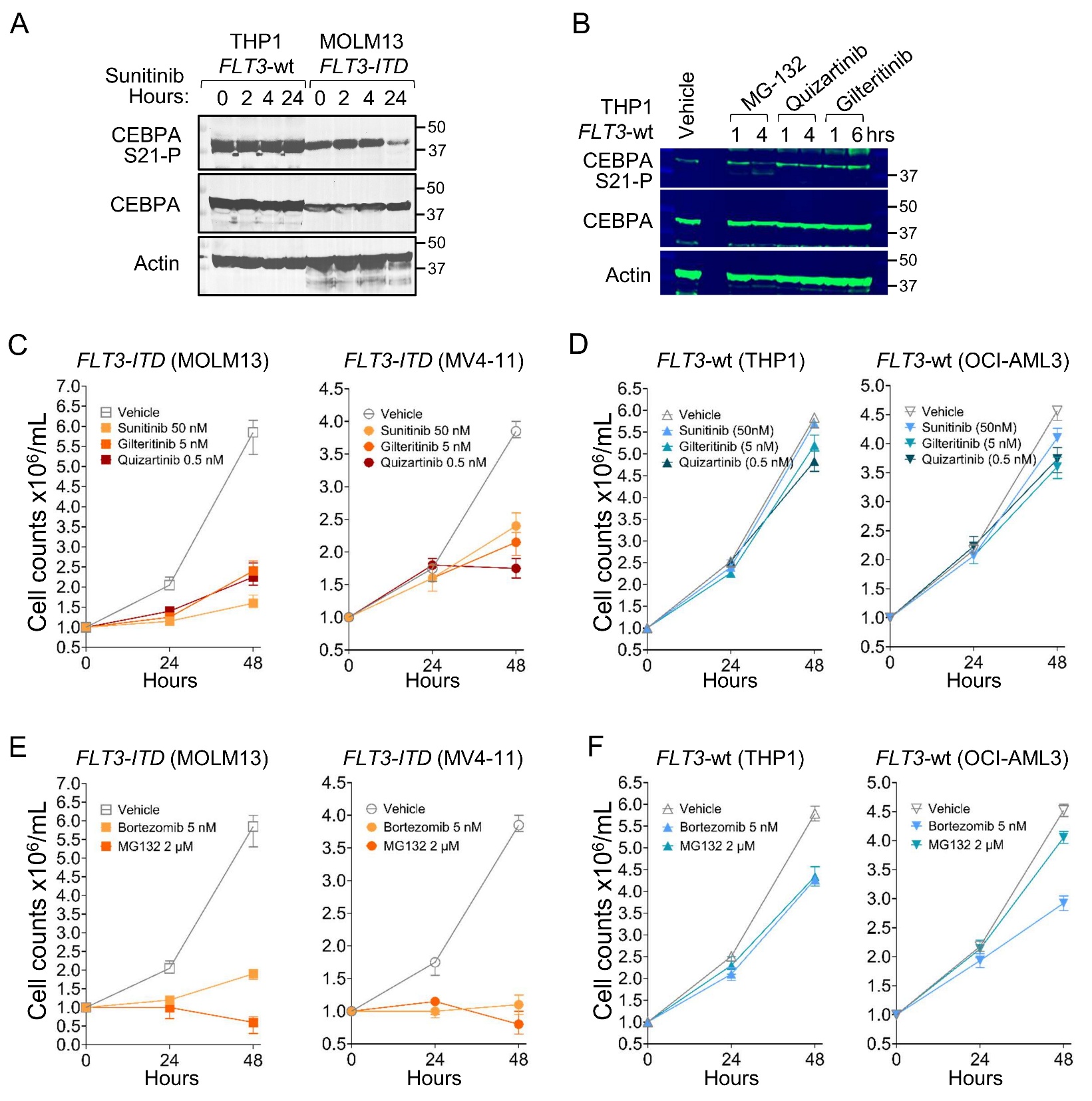

**Figure S3. TKI-treatments did not decrease CEBPA S21-P or increase total CEBPA in *FLT3*-wildtype (wt) AML cells. A) TKI treatment of *FLT3-ITD* AML cells (MOLM13) versus *FLT3*-wt AML cells (THP1).** The cells were treated with vehicle or sunitinib 50 nM and harvested at time-points between 0-24 hours for Western blot analyses. **B) *FLT3*-wt AML cells (THP1) treated with the UPP-inhibitor MG132 500nM, TKI quizartinib 0.5nM or TKI gilteritinib 10nM.** Cells were harvested for Western blots between 1-6 hours after addition of the drugs.

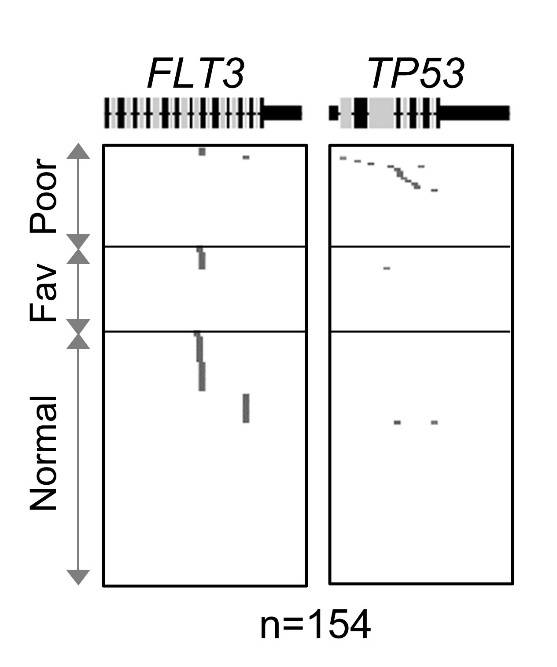

**Figure S4. *FLT3* and *TP53* mutations in AMLs are mutually exclusive, including in the sub-group of AMLs with complex cytogenetics (‘poor’) in which *TP53* mutations are particularly recurrent (The Cancer Genome Atlas [TCGA] public data).**

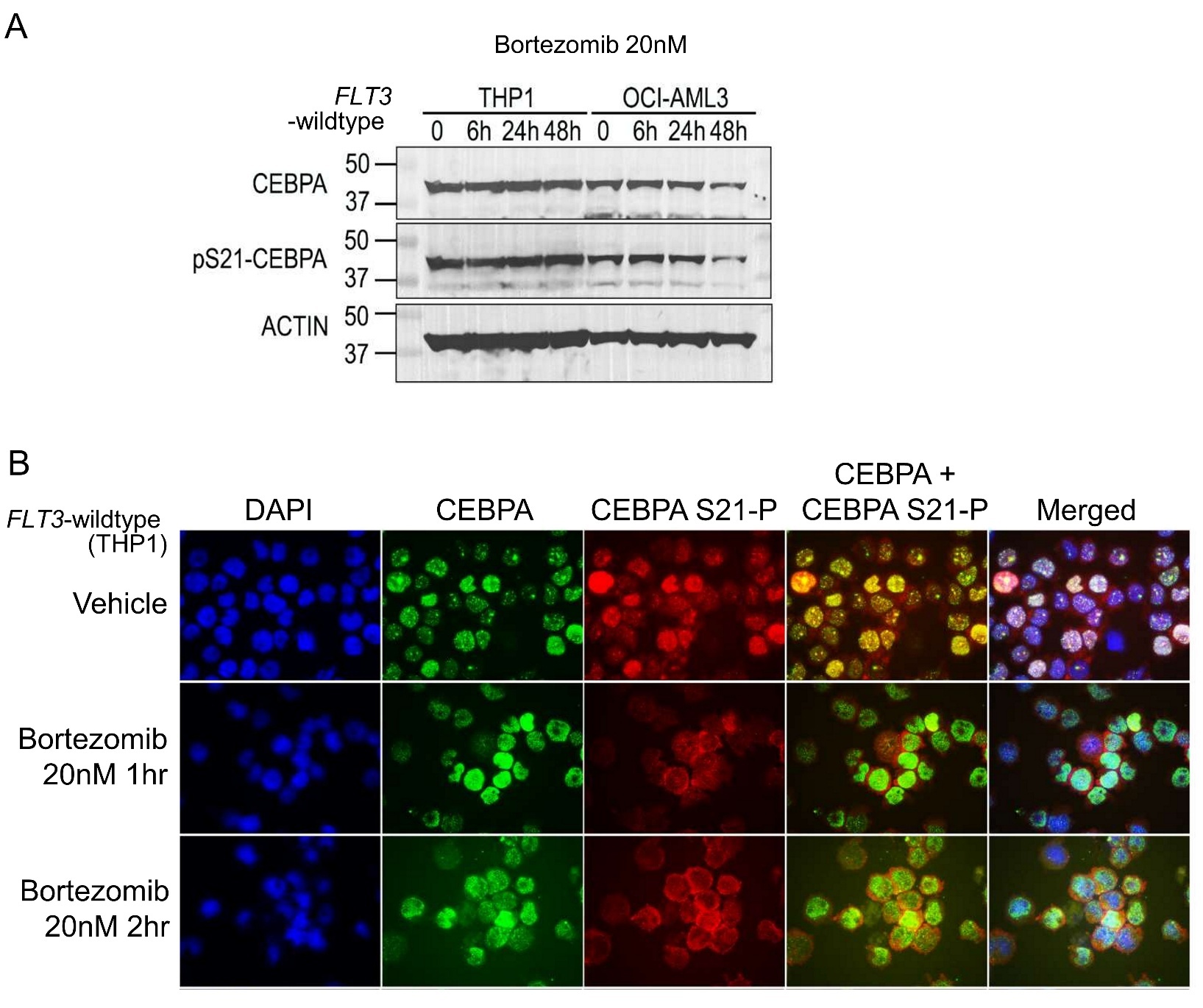

**Figure S5. The UPP-inhibitor bortezomib (UPP-inhibitor MG-132 data in Figure S2) did not increase CEBPA or CEBPA S21-P in *FLT3*-wildtype AML cells THP1 and OCI-AML3**. **A) Western blots in bortezomib-treated THP1 and OCI-AML3 cells.** Concentration and time-points indicated in figure. **B) Western blots in MG132-treated THP1 cells. C) Immunofluorescence for CEBPA and CEBPA S21-P in bortezomib-treated THP1 cells.** Nuclei stained with DAPI. Images by Nikon Eclipse 400 microscope; original magnification, ×630.
